# X-inactivation escapee domains are CTCF-cohesin independent chromatin compartments

**DOI:** 10.64898/2026.08.31.748372

**Authors:** Antonia Hauth, Agnese Loda, Nikolai S. Bykov, Yuvia A. Pérez-Rico, Isabell Rall, Bahtiyar Kurtulmus, Christel Picard, Tim Pollex, Nicolas Servant, Laura Villacorta, Lena Clerquin, Caterina Simoncini, Marc A. Marti-Renom, Edith Heard

## Abstract

X-chromosome inactivation involves chromosome-wide gene silencing accompanied by extensive chromatin changes, as well the loss of topologically associating domains. Yet discrete regions of the inactive X chromosome retain activity within localised 3D domains, which contain active genes that variably escape from X inactivation. The transcription factor and architectural protein CTCF has been proposed to be implicated in escape by insulating escape domains or sustaining their topology via cohesin-mediated loop extrusion. Here, we test the role of CTCF and cohesin in escape using acute degron-mediated depletion of CTCF and RAD21 in neural progenitor cells with established escape profiles. Although CTCF occupancy correlates with escape status on the inactive X chromosome, its removal - together with loss of loop extrusion - does not disrupt escapee gene expression, or domain organization, nor does it result in spreading of silencing or activation of genes in *cis*. Rather, we show that facultative escape regions are self-sustaining compartments of active chromatin enriched in H3K27 acetylation and depleted in H3K27 methylation, with the magnitude of compartment strength scaling up with the degree of transcriptional activity on the inactive X chromosome. These active escapee compartments are propagated independently of CTCF and RAD21-dependent 3D architecture. Our findings identify chromatin compartmentalization as the primary feature of facultative escapee domains.

## Introduction

X-chromosome inactivation (XCI) leads to the transcriptional repression of most X-linked genes on one of the two X chromosomes in female mammals (Loda et al., 2022). The master regulator of XCI is the long non-coding RNA Xist, which becomes monoallelically upregulated during early female development and spreads *in cis* across the X chromosome from which it is transcribed to establish chromosome-wide silencing (Brockdorff et al., 2020). While XCI is essential for female development, some X-linked genes escape silencing and are expressed from both the active (Xa) and inactive (Xi) X chromosomes in female tissues (Carrel and Willard, 2005). Although X-linked escapees are generally expressed at lower levels on the Xi compared to the Xa, their higher expression in females compared to males often results in sex-biased gene expression and can contribute to sex-specific phenotypes in different biological contexts (SB Peeters et al., 2023). For example, variation in XCI escape and escapee gene dosage is increasingly recognized as a key determinant of sex differences in susceptibility to autoimmune, cardiovascular, neurological and metabolic diseases, as well as cancer (Aguado et al., 2022; Davis et al., 2020; Gadek et al., 2025; Hoelzl et al., 2025; Kaneko and Li, 2018; Link et al., 2020; Souyris et al., 2018; Syrett and Anguera, 2019; Youness et al., 2021; Zhang et al., 2024). Despite the growing attention to the role of escapees in sex differences in health and disease, the exact molecular mechanisms that enable this subset of X-linked genes to resist epigenetic silencing by XCI remain largely unknown.

In both humans and mice, a substantial number of genes escape XCI depending on the biological context, accounting for up to 20–25% of X-linked genes (Andergassen et al., 2017; Balaton and Brown, 2016; Berletch et al., 2015; Calabrese et al., 2012; Carrel and Willard, 2005; Cheng et al., 2019; Gendrel et al., 2014; Hauth et al., 2026; Tukiainen et al., 2017). The large majority of escapees evade silencing “facultatively” being transcribed on the Xi only in certain tissues, or developmental contexts, and highly variably across individuals and even cell types (Andergassen et al., 2017; Berletch et al., 2015; Carrel and Willard, 2005; Gadek et al., 2025; Hauth et al., 2026; Hoelzl et al., 2025; Tomofuji et al., 2024; Tukiainen et al., 2017). Previous studies have suggested that escape from XCI is directed by genetically separable cis-regulatory elements, including promoter-proximal dominant elements and distal insulators, although their identity and precise roles remain incompletely defined (Horvath et al., 2013; Li and Carrel, 2008; S Peeters et al., 2023). It has also been proposed that escape may be facilitated by nuclear localisation, as genes that are expressed from the Xi tend to be situated at the periphery, or outside the Xist RNA-coated chromosome during interphase, based on fluorescence in situ hybridization (FISH) experiments (Chaumeil et al., 2006; Chow et al., 2016). Furthermore, whilst escapees generally lack Xist RNA coating (Engreitz et al., 2013; Murakami et al., 2009; Simon et al., 2013), we have recently shown that increased Xist RNA levels can actually override their escape and lead to their silencing. Thus, Xist RNA directly controls the transcriptional activity of escapees in differentiated cells as well as *in vivo*, during development (Hauth et al., 2026). Importantly, transcriptomic and chromosome conformation capture analyses have revealed that variably escaping genes tend to occur in clusters, spanning several hundreds of kilobases and forming 3D topologically associating domain (TAD)-like structures both in embryonic and extraembryonic cells, in contrast to the rest of the Xi where TADs are attenuated (Du et al., 2024; Giorgetti et al., 2016). However, the genesis and function of these domains has remained unknown. It is also unclear whether these TAD-like structures reflect chromatin looping, compartmentalization, or a combination of both. Chromatin loops and compartments result from cohesin-mediated loop extrusion and the segregation of active and inactive chromatin into distinct domains, respectively. Depending on the chromatin context, these mechanisms can either reinforce or antagonize one another (Harris and Rowley, 2024; Hildebrand and Dekker, 2020; Magnitov and de Wit, 2024).

Given that facultative escapees are embedded within these 3D TAD-like domains on the Xi, the multifunctional protein CTCF has been proposed to play a role in their formation and function. As a key architectural factor, CTCF, together with cohesin, may contribute to 3D spatial looping of escapee clusters, thereby facilitating their transcriptional activity on the otherwise silent Xi chromosome territory (Giorgetti et al., 2016). In addition, CTCF has been proposed to insulate escapees and their corresponding domains of active chromatin from neighboring silenced genes and heterochromatin on the Xi (Berletch et al., 2015; Ciavatta et al., 2006; Fang et al., 2025; Filippova et al., 2005; Goto and Kimura, 2009; Horvath et al., 2013). Furthermore, CTCF binds to the promoters of active genes on the Xi where it may directly control their transcriptional activity, in addition to its architectural role (Berletch et al., 2015).

Here we investigate the role of CTCF and RAD21 in escape from XCI, at the X chromosome-wide level using a set of clonal NPC lines that harbor different numbers of facultative escapees, ranging from just a few genes to hundreds. By first mapping the chromatin landscape and structural features of escapee clusters at allelic resolution across multiple clones, and then acutely depleting CTCF or RAD21, we could assess its role on X-linked transcription, chromatin and 3D genome organisation. Unexpectedly, CTCF is found to be largely dispensable for XCI escape in NPCs. Similarly, acute depletion of RAD21 in the same setting had little effect on escape or 3D organisation, revealing that loop extrusion is not the primary driver of 3D topology at escapee clusters. Instead, chromatin states - not looping interactions - govern both the topology and activity of escaping loci on the Xi. Furthermore, no spread of escape into nearby genes, and no spread of silencing into escapees, occurred on the Xi in the absence of CTCF or RAD21. Together these findings resolve a long-standing question by uncoupling CTCF-mediated insulation from escape regulation and uncover a central role for chromatin state in linking 3D architecture to gene expression on the Xi.

## Results

### Transcriptional activity and 3D genome topology at escapee clusters in clonal NPCs

To characterize the transcriptional status and 3D chromatin features of facultative escapees at allelic resolution, we differentiated female F1 hybrid mouse embryonic stem cells (mESC) ((Cast/EiJ) × (C57BL/6)) (Schulz et al., 2014) into NPCs such that random XCI occurs [Fig. 1a]. Starting with single cells taken from a pool of NPCs, we generated a set of clonal cell lines in which either the Cast or the B6 X chromosome is inactive [Fig. 1a]. Allele-specific RNA-seq was performed to determine how many genes show escape from XCI in multiple different clones [Fig. 1b]. We compared the allelic activity status of 407 X-linked genes in ESCs before differentiation, when both X chromosomes are active, and in five independent NPC clones, including two previously characterised clones, E6 and CL30 (Dossin et al., 2020; Hauth et al., 2026). In 3 of the clones (E6, CL30, JTG), the B6 X chromosome is inactive, and in the other two (C5 and B1), the Cast X is inactive. This system allows investigation of chromatin and structural features of Xi escape clusters, independently of the genetic origin of the Xi. In mESCs, most X-linked genes are biallelically expressed prior to XCI and become monoallelically expressed following differentiation and XCI (i.e. their allelic ratio shifts from 0.5 to 0 in NPCs) [Fig. 1b]. Genes that escape XCI result in allelic ratios higher than 0.1 [Fig. 1b]. Our analysis of clonal NPC lines confirmed that, as expected, constitutive escapees are expressed from the Xi in all clones [Fig. 1b]; facultative escapees on the other hand occur in clusters and show considerable variation in escape, ranging from 148 genes escaping XCI in clone E6 to 89 in clone JTG and 84 in clone CL30 [Fig. 1b]. Importantly, this level of variability of escape from XCI, and its mitotic stability, can only be captured when clonal lines derived from single cells are used, highlighting the power of this approach in deconvolving the heterogeneity found at the cell population level (Gendrel et al., 2014; Giorgetti et al., 2016; Hauth et al., 2026).

**Figure 1.**
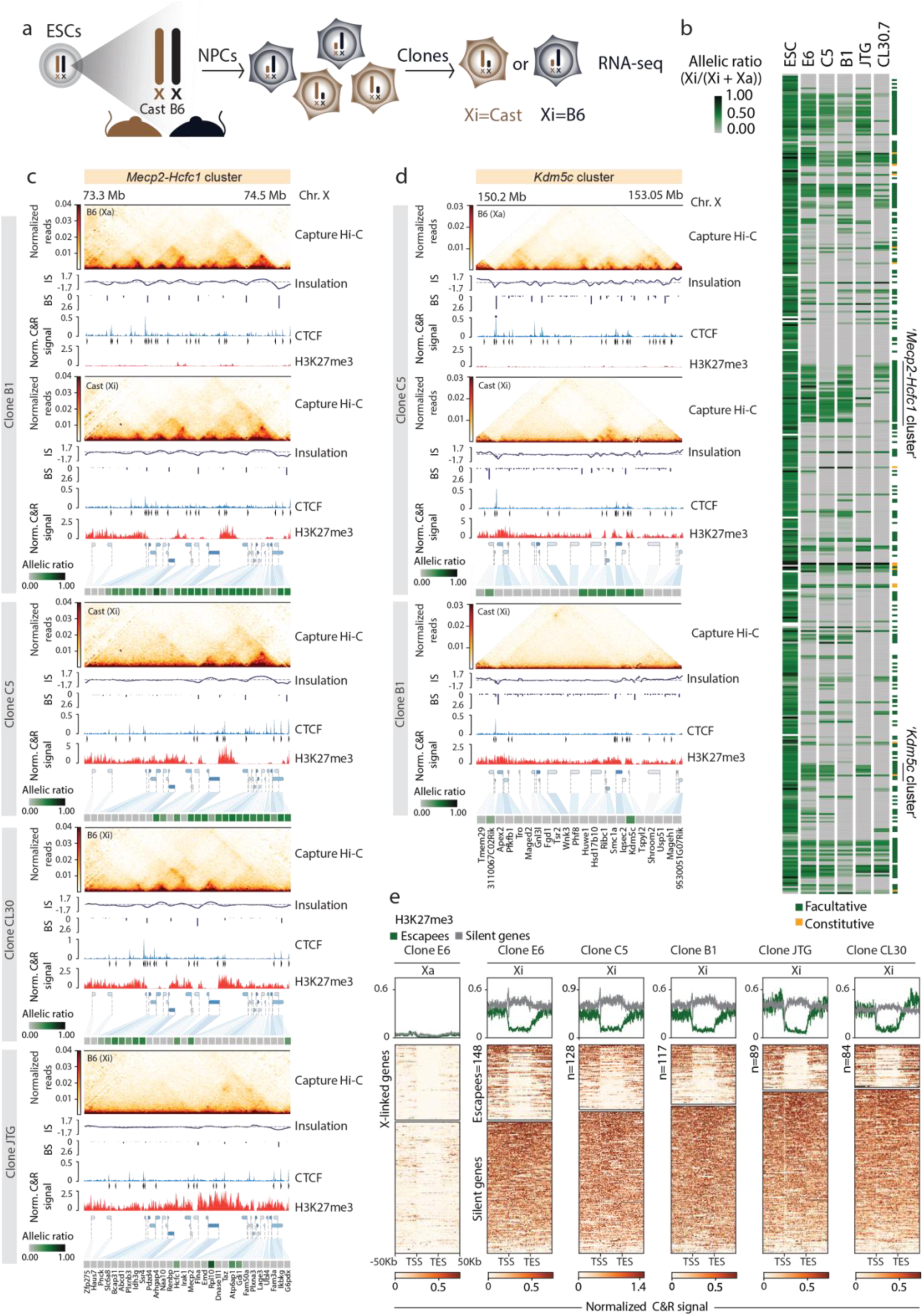
**a**, Experimental workflow for the generation of NPC clones upon differentiation of TX1702 mESCs (C57BL/6J × Cast/EiJ genetic background). Single clones carrying the inactivated B6 or Cast allele were picked and expanded. **b**, Het map showing X-linked allelic ratios (Xi/(Xi+Xa)) in undifferentiated mESCs prior to XCI and in five NPC clones. Allelic ratio indicates the fraction of reads from the Xi compared with reads from the Xi and Xa (ratio,1: Xi monoallelic expression; ratio,0: Xa monoallelic expression; ratio,0.5: biallelic expression; ratio,>0.1: escape). Genes are ordered from centromere (top) to telomere (bottom). Constitutive (yellow) and facultative (green) escapees are indicated. The *Mecp2*–*Hcfc1* and *Kdm5c* escape clusters are highlighted. **c**, Capture Hi-C interaction maps, insulation scores, allele-specific CTCF CUT&RUN profiles and called peaks, and H3K27me3 CUT&RUN profiles at the *Mecp2-Hcfc1* cluster are shown for NPC clones B1, C5, CL30 and JTG. The Xa is shown for clone B1. Capture Hi-C data are shown at 10 kb resolution. CTCF motif orientation is indicated by arrowheads. Heatmaps of allelic ratios for genes within the cluster are shown.. **d**, As in **c**, for the *Kdm5c* escape cluster in NPC clones C5 and B1. **e**, Aggregate profiles (top) and heat maps (bottom) of H3K27me3 signal across X-linked escapee (green) and silent (grey) genes are shown. The Xa is shown for clone E6. The Xi is shown for clones E6, C5, B1, JTG and CL30. RNA-seq and CUT&RUN data represent two biological replicates; Capture Hi-C data are from a single replicate. [IS - insulation score; BS - boundary score]

To investigate the 3D organization of facultative escapee clusters on the Xi, we focused on two large regions, one of which spans 1 Mb and includes the *Mecp2-Hcfc1* gene cluster, while the other spans 3 Mb and includes the cluster with the *Kdm5c* gene. These two regions have previously been shown to form TAD-like structures on the Xi when genes within them display escape (Giorgetti et al., 2016; Hauth et al., 2026). We resolved the 3D topology of these two regions at 10 kb resolution by performing allele-specific Capture Hi-C analysis in NPC clones exhibiting different levels of escape [Fig. 1c, Extended Data Fig. 1a-c].

Within the *Mecp2-Hcfc1* cluster, the numbers of facultative escapees ranged from 26 in B1 to 18 in C5, 10 in CL30, and 4 in JTG [Fig. 1c]. We observed clone-specific differences in the 3D organization of this cluster on the Xi, which largely reflect the transcriptional activity within the region [Fig. 1c]. On the active Xa no differences in 3D organisation of this region were detected across all clones analyzed [Extended Data Fig. 1a]. In clone B1, where the entire cluster of genes escape from XCI, the 3D organization on the Xi closely resembles that of the Xa; in contrast, in clone JTG, where only four genes escape XCI, this region on the Xi lacks clear TAD-like organization [Fig. 1c]. These features are consistent in clone C5 and CL30 as well, with the 3D topology of the cluster reflecting different expression states of escapees [Fig. 1c].

Similar observations were made for the *Kdm5c* cluster, with a TAD-like organization present on the Xi in clone C5 where 8 genes are active. In clone B1 where only two genes escape XCI in this region, no domain is apparent and only a single long-range loop is detected [Fig. 1d]. On the active X, the topology of the *Kdm5c* region was the same in all clones analysed, similarly to the case for the *Mecp2-Hcfc1* cluster [Extended Data Fig. 1b].

We also assessed CTCF binding and H3K27me3 enrichment within these regions using allele-specific CUT&RUN analysis [Fig. 1c-d]. H3K27me3 enrichment is normally found across the majority of the inactive X chromosome. Within the *Mecp2-Hcfc1* and *Kdm5c* clusters, a distinct lack of H3K27me3 deposition is found over the gene bodies of escapees when they are actively transcribed on the Xi.

This contrasts with H3K27me3 enrichment across silent genes and leads to distinctive H3K27me3 patterns within escapee cluster regions amongst the different clones [Fig. 1c-d]. Such a depletion of H3K27me3 across gene bodies, from transcription start sites (TSS) to transcription end sites (TES), of active escapees, but not of silent genes, was observed chromosome-wide across all escape regions of the Xi [Fig. 1e]. Again, distinctive patterns of H3K27me3 distribution could be seen amongst clones exhibiting varying levels of escape [Fig. 1e].

Visualization of CTCF binding across the *Mecp2-Hcfc1* and *Kdm5c* clusters confirmed globally reduced CTCF occupancy on the Xi relative to the Xa [Fig. 1c–d, Extended Data Fig. 1a-c] (Berletch et al., 2015; Du et al., 2024; Minajigi et al., 2015), except at escape loci within the clusters where CTCF was still present [Fig. 1c–d]. Furthermore, clone-specific profiles of retention of CTCF aligned well with regions of active transcription within escapee clusters on the Xi [Fig. 1c–d], as might be expected if CTCF plays a role in permitting escape.

In summary, our integrated profiling of allelic gene expression, CTCF occupancy, H3K27me3 enrichment, and 3D chromatin architecture across five independent NPC clones reveals that facultative escape clusters adopt clone-specific chromatin and 3D topological states that closely mirror their transcriptional activity on the Xi. This also highlights the substantial variability in facultative escape status in genetically identical clones.

### Escapees retain CTCF binding and H3K27ac enrichment according to XCI status

We next investigated whether similar allele- and clone-specific CTCF binding patterns observed at the *Mecp2-Hcfc1* and *Kdm5c* clusters, could be identified at other escapee gene clusters on the Xi. We explored the genome-wide CTCF binding patterns by first detecting peaks on the total signal in each clone [Extended Data Fig. 2a]. For each detected peak, we calculated an allelic ratio similarly to the strategy used for allele-specific RNA-seq analysis (C&R allelic ratio = (Xi/(Xi+Xa); ratio >0.25 biallelic peaks; ratio <0.25 Xa-monoallelic peaks). Comparing allelic ratios of CTCF peaks on the X chromosome to those on autosomes revealed that, while autosomal peaks are largely biallelic, CTCF binding on the X chromosome is skewed toward the Xa, confirming higher occupancy on the Xa relative to the Xi [Extended Data Fig. 2b] (Berletch et al., 2015; Minajigi et al., 2015). The Xi escapees in all five NPC clones revealed higher CTCF binding at their TSS compared to those of silenced genes on the Xi, thus extending our observation at the *Mecp2-Hcfc1* and *Kdm5c* clusters to the entire Xi [Fig. 2a]. We also performed comparisons of the allelic ratios of CTCF peaks at promoters and gene bodies of escapees between clone pairs. This analysis showed that CTCF binding at escapees is maintained on the Xi only when the gene is active and lost when the same gene is silent in a different clone [Fig. 2b-c].

**Figure 2.**
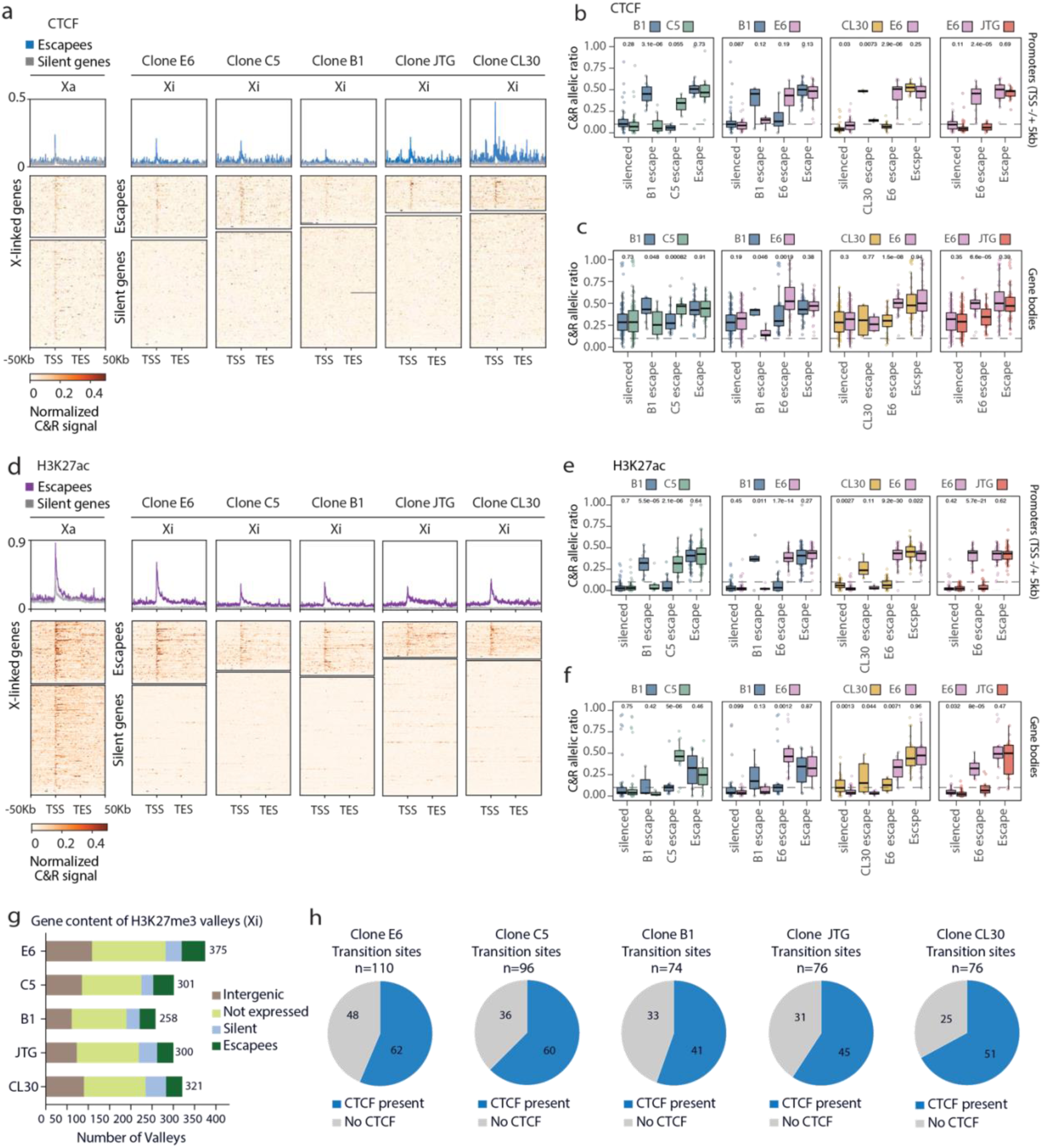
**a**, Aggregate profiles (top) and heat maps (bottom) CTCF C&R signal on the Xi across gene bodies and ±50 kb flanking regions of X-linked escapees (blue) and silent genes (grey). The Xa is shown for clone E6. The Xi is shown for clones E6, C5, B1, JTG and CL30. **b**, Boxplot showing pairwise comparison of allelic ratios of promoter-associated CTCF peaks (TSS ±5 kb) between NPC clone pairs, stratified by escape category (silenced in both clones, escaping in both clones, or escaping in one clone only). **c**, As in **b**, but for gene body-associated peaks. **d-f**, As in **a-c**, for H3K27ac peaks. Only groups with >2 data points are shown. **g**, Gene content of H3K27me3 valleys on the Xi in NPC clones E6, C5, B1, JTG and CL30. Valleys were identified from clone-specific, replicate-merged, normalised H3K27me3 CUT&RUN bigWig tracks at 5 kb resolution using a two-state Gaussian Hidden Markov Model (hmmlearn v0.3.3, GaussianHMM, n_components = 2, covariance_type = “diag”), with ENCODE mm10 blacklist v2 regions excluded from the training set. Stacked bars show the total number of valleys per clone stratified by genomic feature and gene content (intergenic, not expressed, silent X-linked genes, escapees); categories were assigned by intersection with the gene annotation and the clone-specific RNA-Seq allelic ratio classification. 12% of H3K27me3 valleys overlap with silent genes, 45% with not expressed genes, 28% with intergenic regions, and 15% with transcriptionally active escapees. **h**, CTCF enrichment at active–inactive chromatin transition sites flanking H3K27me3 valleys containing at least one escapee, shown as pie charts for each NPC clone. Sites were classified as “CTCF present” (blue) if a clone-specific CTCF CUT&RUN peak was located within 50 kb inward and 10 kb outward of the transition site, or as “No CTCF” (grey) otherwise (see methods). *n*, total number of transition sites analyzed per clone.

We then profiled the enrichment of H3K27ac, one of the histone marks that is rapidly lost at promoters of X-linked genes that become silenced during XCI but retained at loci encompassing escapees (Toothacre et al., 2026; Żylicz et al., 2019). We performed this analysis in five clones and found that the majority of H3K27ac peaks detected on the X chromosome and on autosomes are shared in at least two clones [Extended Data Fig. 2c]. Similar to CTCF, H3K27ac peaks at the X chromosome-wide level are largely biased toward the Xa, highlighting the general lack of active chromatin on the Xi, where most X-linked genes are inactive in NPCs [Extended Data Fig. 2d]. At loci encompassing active escapees, H3K27ac enrichment resembles the pattern observed for CTCF, both at escapees TSS and in pairwise comparisons of promoter and gene body enrichment between clones [Fig. 2d-f]. Thus, in NPCs, enrichment of both CTCF binding and H3K27ac at X-linked genes on the Xi directly reflects the number and degree of escapees across different clones [Fig. 2a-f].

Next we used our genome-wide CUT&RUN data to define active and inactive chromatin domains on the Xi. In this way we could examine CTCF occupancy at transition sites between active/inactive chromatin, given that CTCF has been proposed to act as a potential insulating factor for escapee regions [Berletech 2015, Filippova 2005, Goto 2009, Horwarth 2013, Ciavatta 2006, Fang 2025]. To define transition sites between active and inactive chromatin domains across all 5 NPC clones we integrated RNA-seq data with CUT&RUN for H3K27me3 and H3K27ac [Fig. 2g, Extended Data Fig. 2e-g]. First, using a 2-state Hidden Markov Model (HMM) we determined which regions on the Xi show a significant decrease in H3K27me3 enrichment compared to their neighbouring regions. We focused on H3K27me3 because, within both the *Mecp2-Hcfc1* and *Kdm5c* clusters, we observed discrete domains depleted of H3K27me3 at transcriptionally active regions, with sharp boundaries, and these varied between clones according to escapee activity, consistent with clone-specific patterns of XCI escape [Fig. 1c,d]. This approach delineated “valleys” of H3K27me3 on the Xi, for which numbers vary across clones [Fig. 2g].

Notably, this X chromosome-wide analysis revealed numerous H3K27me3 valleys that not only overlap with escapees but also with silenced genes, non-expressed regions, or gene deserts [Fig. 2g]. These observations suggest that local depletion of H3K27me3 alone is likely not sufficient to define transition sites between active and inactive chromatin on the Xi. To further investigate the features that distinguish escape domains, we examined which H3K27me3 valleys were enriched for H3K27ac and found that escapee-containing valleys display significantly higher levels of H3K27ac compared with all other valleys [Extended Data Fig. 2e-f]. Thus, it is not simply the absence of H3K27me3, but rather the combination of H3K27me3 depletion and H3K27ac enrichment that characterizes the transition sites between active and inactive chromatin domains on the inactive X chromosome [Extended Data Fig. 2e-f].

Finally, to assess CTCF binding at these transition sites, we analyzed regions extending 10 kb outward and 50 kb inward from each edge of escapee valleys and observed CTCF occupancy at 55% (clone B1) to 67% (clone CL30) of these sites, with similar levels across clones [Fig. 2h, see methods]. These results indicate that CTCF is enriched at a subset, but not at all transition sites separating silent and active domains on the Xi [Fig. 2h]. Nevertheless, when we compared CTCF peaks at the edges of valleys encompassing escapees, with those associated with gene deserts and silent genes, we found that CTCF occupancy was significantly associated with escape status, indicating that its enrichment at escape valleys is non-random (Fisher’s exact test, p < 1 × 10⁻¹³) [Extended Data Fig.2g].

These results show that most transition sites between active and inactive domains on the Xi are enriched with CTCF. Furthermore CTCF localizes at active escapees on the Xi, where its enrichment overlaps with H3K27ac. Together, these results suggest that CTCF may regulate escapees at multiple levels.

### Compartment strengthening and loop maintenance at Xi escape domains

We next set out to define how chromatin looping and compartmentalization within escapee domains differ between the Xa and Xi, in particular between different clones with varying levels of XCI escape [Fig. 3a-c]. In our previous work we discovered TAD-like structures on the Xi at 40 kb resolution (Giorgetti et al., 2016). Here we increased resolution thanks to our Capture Hi-C data at the *Mecp2-Hcfc1* and *Kdm5c* clusters, that achieves 10 kb resolution. This allowed us to investigate the interplay between looping and compartmentalization within escapee domains on the Xi. We first ranked NPC clones according to their level of escape based on RNA-seq by defining an average cluster D-score for each clone, calculated across all genes within the *Kdm5c* and *Mecp2-Hcfc1* clusters [Fig. 3a,b]. The average cluster D-score corresponds to the mean allelic ratio (Xi/(Xa+Xi)) computed across all expressed genes within a given cluster. By integrating both the number of escapees and their relative expression on the Xi, this provides a continuous, cluster-level readout of escape that allows direct comparison between clones with otherwise heterogeneous escape patterns (see methods). The average cluster D-scores at the *Mecp2*-*Hcfc1* cluster ranged from 0.05 in clone JTG, in which only three genes escape XCI [Fig. 1c], through to 0.41 in clone E6, in which the majority of genes within the cluster are expressed from the Xi [Fig. 3b]; a similar gradient was observed at the *Kdm5c* cluster, with average cluster D-scores ranging from 0.04 (B1) to 0.20 (E6) [Fig. 3a].

**Figure 3.**
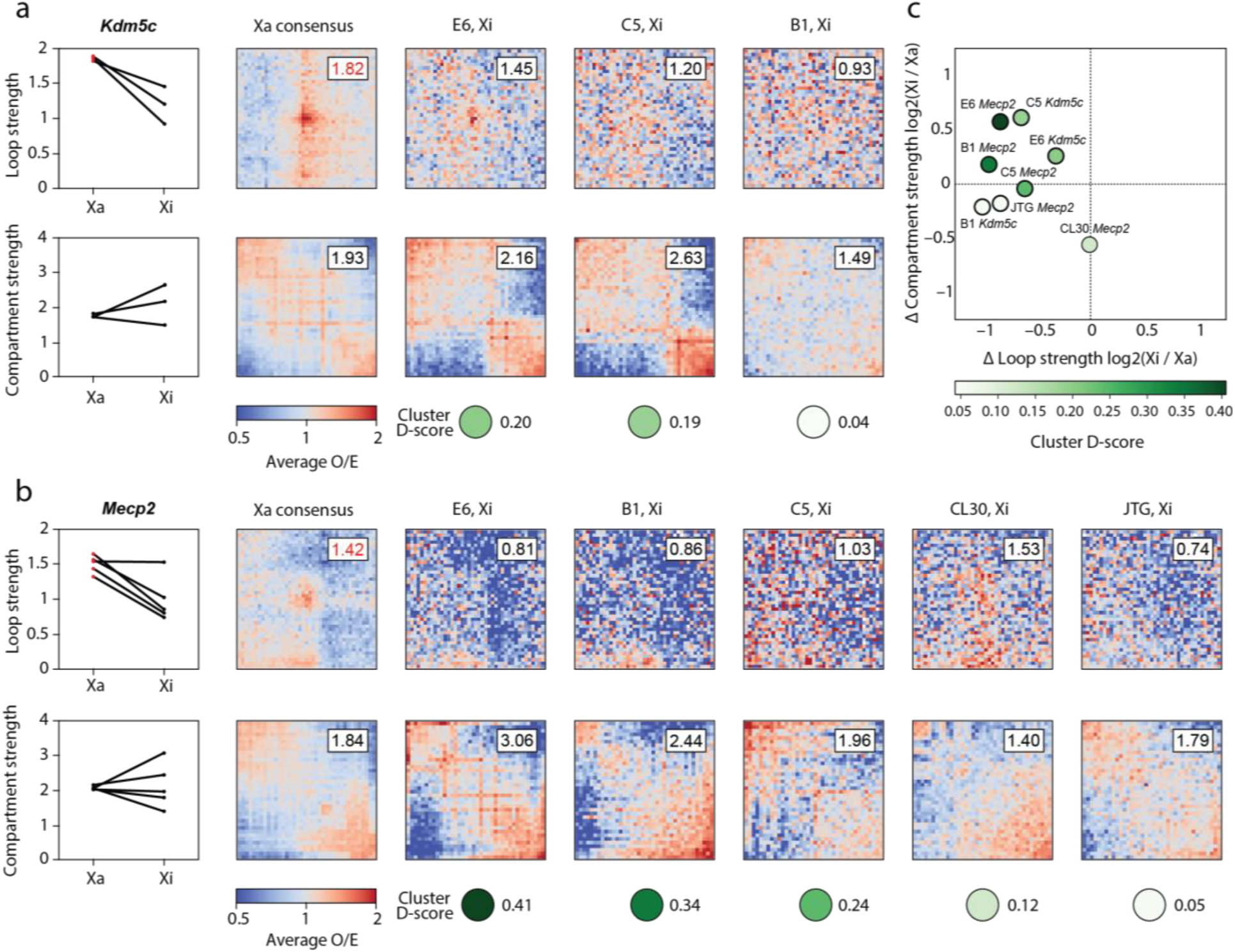
**a**, Loop and compartment strength at the *Kdm5c* cluster across NPC clones E6, C5 and B1. Top: per-clone mean loop strength on Xa and Xi (one line per clone; loops were called on Xa), together with average observed-over-expected loop pile-up matrices at 5 kb resolution centred on consensus *Kdm5c* loops (see methods); mean O/E in the central 3 × 3 pixels is shown at top right of each pile-up. The Xa consensus pile-up uses the merged Xa cooler (see methods); Xi pile-ups are clone-specific (E6, C5, B1). Green circles below each pile-up reflect the per-clone average cluster D-score. Bottom: compartment strength. Saddle plots are shown for the Xa consensus (see methods) and clone-specific Xi chromosomes **b**, As in **a** for the *Mecp2-Hcfc1* cluster, across NPC clones E6, B1, C5, CL30 and JTG (Xi) and the merged Xa consensus (see methods). **c**, Scatter plot of compartment-strength change versus loop-strength change (Xi relative to Xa) for both loci across all NPC clones. Each dot represents one clone × locus, positioned at x = log2(Xi/Xa) loop strength and y = log2(Xi/Xa) compartment strength, and coloured by the average cluster D-score of that clone × locus.

Next, we compared chromatin loop strength between the Xa and the Xi chromosomes across clones with different levels of escape from XCI. To define a common set of loops for comparison, we first identified focal interactions on the Xa using Capture Hi-C maps merged across all clones generated in this study (see Methods). We then performed allele-specific pile-up analysis for each clone at these consensus loop coordinates. Loop strength was quantified as the mean observed/expected contact enrichment at the loop centre relative to the surrounding local background [Fig. 3a–b; see methods].

At the *Kdm5c* cluster, loops are attenuated on the Xi in clones E6, C5 and B1 when compared to the same region on the Xa [Fig. 3a,c]. Comparison of loop strength on the Xi in the three clones showed that higher escape numbers and levels are associated with stronger looping interactions on the Xi [Fig. 3a,c]. At the *Mecp2-Hcfc1* cluster we also found a general decrease in loop strength on the Xi compared to the Xa across clones [Fig. 3b] (with the exception of clone CL30). However, at this cluster we did not find a clear correlation between loop strength and escape levels when comparing the Xi in clones ranked by degree of escape levels [Fig. 3b,c]. Thus, increased escape levels correlate with stronger loops in some but not all cases.

We next compared compartment strength at the *Kdm5c* and *Mecp2-Hcfc1* clusters by applying a saddle-plot analysis [Fig. 3a,b]. Bins were ordered along the principal component eigenvector computed from the Capture Hi-C maps, and compartment strength was defined as the ratio of average homotypic (AA + BB) to heterotypic (AB + BA) interactions [Fig. 3a-b, see methods]. Compartmentalization on the Xi was notably reinforced with increasing escape, with compartment strength being generally positively associated with escape levels. This trend was consistently observed for the *Kdm5c* cluster (3/3 clones) and the majority of clones at the *Mecp2-Hcfc1* cluster (4/5 clones) [Fig. 3b,c].

To determine the statistical significance of the observed loop-versus-compartment trends at both the *Mecp2*-*Hcfc1* and *Kdm5c* clusters [Fig. 3c] and obtain an overall view of these features beyond any single locus, we applied the same framework to these regions on their Xa alleles [Extended Data Fig. 3a-b, see methods]. Mean loop strength was computed for the Xi and Xa within these regions [Extended Data Figs 3a]. Compartment strength was also computed independently on the Xa and Xi at matching coordinates [Extended Data Fig. 3b]. Having a panel of multiple clones at the same loci enabled us to assess statistical significance of the difference between Xa and Xi for each of these two features (i.e. loop extrusion and compartmentalisation) [Extended Data Fig. 3, see methods]. Across clones, mean loop strength was significantly higher on the Xa than on the Xi (Wilcoxon signed-rank test, one-sided Xa > Xi, p = 3.1 × 10⁻⁴) [Extended Data Fig. 3a]. In direct contrast, mean compartment strength was significantly lower on the Xa than on the Xi over the same clones (one-sided Xa < Xi, p = 0.037) [Extended Data Fig. 3b]. Thus, our data indicate that on the Xi the attenuation of CTCF/cohesin-dependent looping is accompanied, and likely opposed by, a reinforcement of compartmentalisation [Fig. 3c], in line with the previously described antagonism between loop extrusion and compartmentalisation as drivers of 3D genome organisation (Nuebler et al., 2018).

This antagonism between looping and compartmentalisation may also explain the different relationships between looping strength and escape levels observed at the *Kdm5c* and *Mecp2-Hcfc1* clusters [Fig. 3c]. Although escape at both clusters generally occurs within the context of reduced looping and enhanced compartmentalization relative to the Xa, the balance between these two features may differ locally, such that escape at the *Kdm5c* cluster is more closely associated with looping, whereas compartmentalization may represent the dominant structural feature at the *Mecp2-Hcfc1* cluster [Fig. 3c].

In summary, our data shows that both chromatin looping and compartmentalization contribute to the organization of escapee domains on the Xi, potentially through independent and counterposing mechanisms.

### CTCF is dispensable for escape maintenance on the Xi

Given the potential roles that CTCF may have in regulating escape, we established a CTCF dTAG-based degron system, which allows acute depletion of CTCF protein upon the addition of the small molecule dTAG to the culture media (Nabet et al., 2018) [Fig. 4a]. We generated eight CTCF-dTAG NPC clones in which the FKBP12^F36V^ degron tag and the fluorescent reporter GFP are integrated at the C terminus of endogenous CTCF on both alleles [Fig. 4b]. This set of NPC clones includes four clones generated by direct CRISPR-Cas9 mediated targeting of NPC clones B1, C5 and E6 (i.e. B1D9, B1G1G3, C5C10, E6A7), and four clones derived upon differentiation of TX1072 ESCs, in which the FKBP12^F36V^ tag had been targeted to the CTCF endogenous locus in ESCs before cell differentiation [Fig. 4a-c]. NPC clones B1, C5, and E6 retained their original escape profiles and expression levels following the CTCF-tagging process. This highlights the stability of escape status of X-linked genes, which once established is epigenetically maintained not only through cell proliferation, but also upon NPC subcloning [Extended Data Fig. 4a]. The four NPC clones that were derived from targeted ESCs showed slightly lower levels of escape, both in terms of number of genes escaping XCI and their expression levels [Fig. 4c]. This is probably because they were derived under conditions of doxycycline-induced overexpression of Xist on the BL6 X chromosome during differentiation, which results in reduced facultative escape, as previously reported (Hauth et al., 2026).

**Figure 4.**
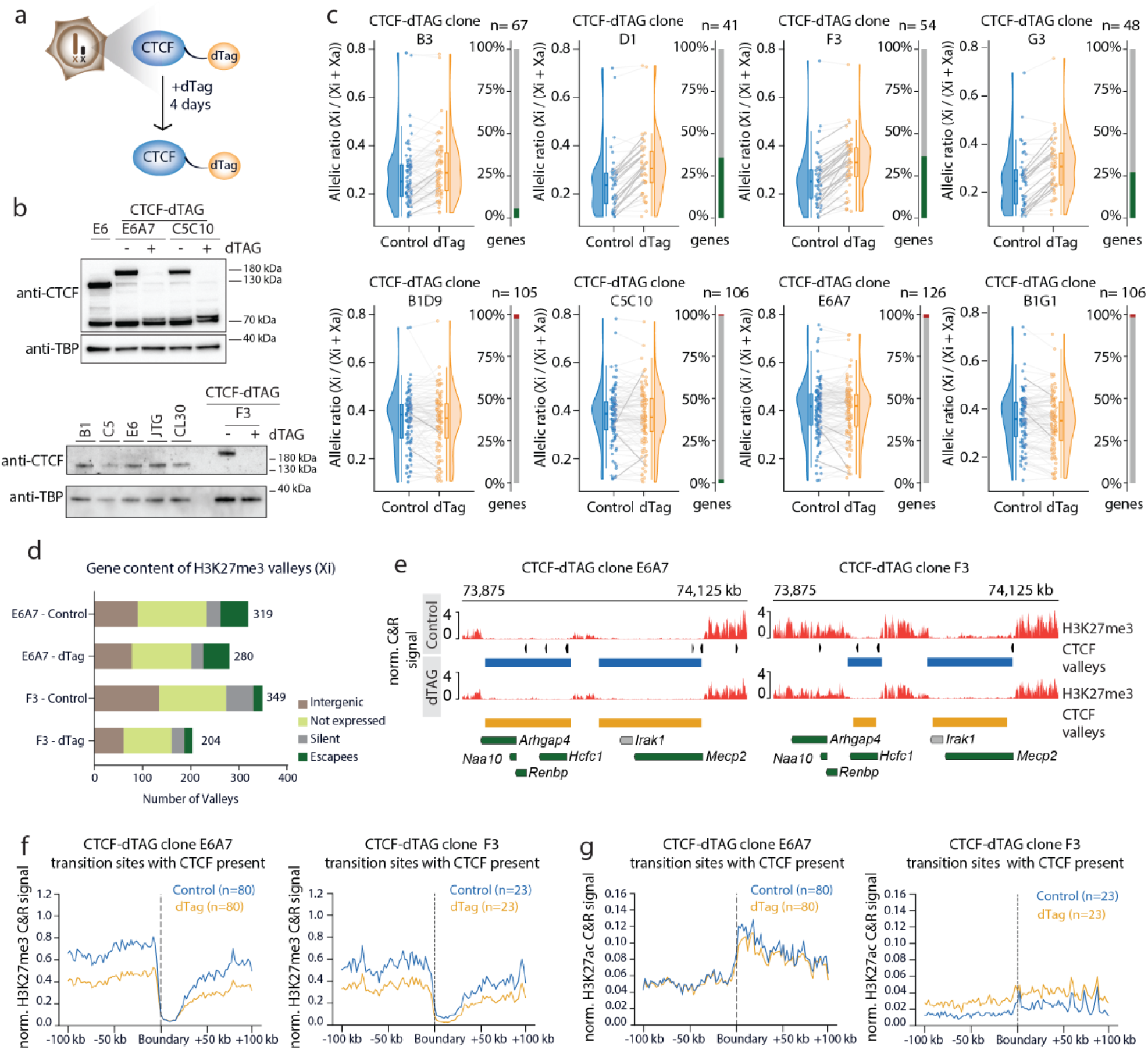
**a**, Schematic of the experimental design for acute CTCF degradation in NPCs. **b**, Western blots showing CTCF protein levels in CTCF-dTAG clones (E6A7, C5C10 and F3) treated (+) or untreated (−) with dTAG. NPC clones B1, C5, E6, JTG and CL30 are shown for comparison. TBP served as the loading control. **c**, Raincloud plots showing allelic ratios for X-linked genes with allelic ratio > 0.1 in either control or dTAG-treated condition across CTCF-dTAG NPC clones. Plots show the density distribution, boxplot, and paired allelic ratio for individual genes in control and dTAG-treated conditions. Bar plots indicate the percentage of genes exhibiting a significant increase (green) or decrease (red) in allelic ratio following dTAG treatment (Δallelic ratio > 0.05, adjusted *P* < 0.05). A small number of escapees showed a significant decrease in allelic ratio upon CTCF degradation: *Gpm6b*, *Gprasp1* and *Syp* in clone B1D9; *Eif1ax* in clone C5C10; *Alg13*, *Eif1ax* and *Gprasp1* in clone E6A7; and *Gpm6b* and *Gprasp1* in clone B1G1. *Fam199x* also showed a significant decrease in allelic ratio in clone C5C10 and fell below the allelic ratio threshold of 0.1 following dTAG treatment; therefore, it is not included in the plot. **d**, Gene content of H3K27me3 valleys on the Xi in CTCT-dTAG clones E6A7 and F3. Valleys were identified as in Fig. 2i. **e**, Genome browser views of H3K27me3 CUT&RUN signal across part of the *Mecp2–Hcfc1* cluster on the Xi in CTCF-dTAG clones E6A7 (left) and F3 (right), in the control condition (top) and following dTAG treatment (bottom). CTCF peaks are indicated by arrowheads pointing in the motif direction. H3K27me3 valleys are shown for each condition. Escapees (green) and silent genes (grey) are shown. **f**, H3K27me3 signal at CTCF-associated active–inactive chromatin transition sites on the Xi in CTCF-dTAG clones E6A7 (left) and F3 (right). Mean signal from normalised, replicate-merged CUT&RUN bigWig tracks at 5 kb resolution was aggregated across 100 kb windows centred on transition sites associated with a clone-specific CTCF CUT&RUN peak (located within 50 kb inward and 10 kb outward of the transition site) for control (blue) and dTAG-treated (orange) conditions. n, number of transition sites. Normalisation was performed with csaw v1.42.0/edgeR-TMM as described in the methods. **g**, as in **f** for H3K27ac.

Next, we depleted CTCF for 4 days in all eight NPC clones. Efficient CTCF degradation upon dTAG treatment was confirmed by Western Blot [Fig. 4b] or by flow cytometric analysis of CTCF-GFP levels [Extended Data Fig. 4b]. CTCF CUT&RUN analysis before and after dTAG treatment further confirmed robust CTCF depletion at the chromatin level on the X chromosomes and autosomes [Extended Data Fig. 4c-d]. Genome-wide RNA-seq of up- and down-regulated genes revealed overall similarities with some differences between clones following CTCF depletion [Extended Data Fig. 4e].

We then assessed changes in allelic ratios of X-linked escapees upon CTCF loss [Fig. 4c]. If CTCF is directly involved in ensuring the transcriptional activity of escapees, its depletion should lead to allelic ratios falling below the 0.1 threshold set for XCI escape. However, no major changes in allelic ratios of escapees were observed. In fact, we even detected a slight increase in expression for some escapees with lower basal expression [Fig. 4c]. In the case of silent genes (i.e. allelic ratio <0.1), out of 361 analyzed genes, only 12 showed increased allelic ratio above the 0.1 threshold used to call escapees [Extended Data Fig. 4f]. Notably, all 12 genes have previously been reported to escape XCI in NPCs (Hauth et al., 2026) and displayed low levels of Xi expression in the absence of dTAG treatment, with allelic ratios greater than 0 but just below the threshold used to define escape prior to CTCF depletion [Extended Data Fig. 4f] This data demonstrates that CTCF depletion does not lead to broad reactivation of silent genes on the Xi.

We examined whether the small changes in allelic ratios of some escapees upon loss of CTCF were due to a change in transcriptional activity on the Xi and/or the Xa [Extended Data Fig. 5a]. We performed differential expression analysis of escapees on the Xa and Xi independently, in dTAG versus control conditions [Extended Data Fig. 5a]. This analysis revealed that the escapees displaying increased allelic ratios upon CTCF loss were significantly upregulated on the Xi, but not on the Xa [Extended Data Fig. 5a]. We also investigated whether this relative upregulation might be the result of indirect changes in Xist RNA levels upon CTCF loss, as we previously showed that Xist RNA directly regulates escapee expression levels in NPCs (Hauth et al., 2026). CTCF depletion did not lead to decreased Xist levels; in fact, a modest increase was observed in most NPC clones, arguing against an indirect effect mediated by Xist RNA levels in the absence of CTCF [Extended Data Fig. 5b]. In the four NPC clones with higher baseline levels of escape, our differential expression analysis upon dTAG treatment showed largely unchanged escapee expression levels upon CTCF loss on both the Xa and Xi [Extended Data Fig. 5c]. Thus, escapees expressed at lower levels in some NPC clones seem to retain a greater tendency for relative upregulation upon CTCF loss, whereas changes in the same genes, if initially expressed more highly, are unaffected or undetectable following CTCF loss.

Overall these results show that even though CTCF is enriched at escapees and domains, it is in fact largely dispensable for the maintenance of escape in NPCs. Acute depletion across eight independent NPC clones did not alter the escape status of escapees on the Xi, it actually resulted in a modest increase in expression, in some of the NPC clones, and silent X-linked genes remained repressed in the absence of CTCF.

### Chromatin state transitions on the Xi are maintained independently of CTCF in NPCs

Next, we assessed the role of CTCF as an insulator of escapee regions. We thus tested whether loss of CTCF alters the organization of active and inactive chromatin domains on the Xi, and whether such chromatin reorganization leads to spreading of escape-associated chromatin domains into silenced regions of the Xi. First, we determined whether CTCF loss alters the number and extent of H3K27me3 valleys on the entire Xi [Fig. 4d]. We performed this analysis in clones E6A7 and F3. These two clones were selected as they show high (E6A7) and low (F3) escape levels at the X-chromosome-wide levels and within the *Mecp2-Hcfc1* and *Kdm5c* clusters. We found that CTCF depletion does not dramatically affect escapee valleys but leads to a general decrease in numbers of H3K27me3 valleys encompassing intergenic regions, gene deserts and not expressed genes across the Xi, particularly in clone F3 [Fig. 4d]. These overall changes in H3K27me3 valleys likely reflect the fact that CTCF loss leads to a global reduction in H3K27me3 levels, not only on the Xi, but also on autosomes, consistent with previous reports [Fig. 4d, Extended Data Fig. 6a] (Nora et al., 2017). To quantify the effect of CTCF depletion on escape-associated valleys, we specifically examined H3K27me3 valleys encompassing escapees, the majority of which are bound by CTCF at their edges [Extended Data Fig. 6b]. We found that these transition sites between escapee domains and silent regions of the Xi remain largely unchanged upon CTCF loss, with 80% and 67% of valleys detected prior to dTAG treatment overlapping those identified following CTCF depletion in clones E6A7 and F3, respectively [Fig. 4d; Extended Data Fig. 6c].

Although escapee valleys persisted in the absence of CTCF, indicating neither spreading of H3K27me3 into active escapee loci, nor enlargement of H3K27me3-depleted regions upon CTCF loss, we detected reduced H3K27me3 levels surrounding their edges, consistent with the observed genome-wide decrease in H3K27me3 [Fig. 4e-f]. These observations demonstrate that CTCF is largely dispensable for maintenance of the separation between active and inactive domains on the Xi [Fig. 4e-f]. This is in line with the RNA-seq analysis showing no reactivation of silenced X-linked genes and no silencing of escapees within H3K27me3 valleys following CTCF depletion [Extended Data Fig. 4f]. Finally, we examined H3K27ac enrichment within escapee valleys following CTCF loss [Fig. 4g]. In clone E6A7, H3K27ac levels remained unchanged, in agreement with the absence of significant changes in escapee expression upon CTCF depletion in this clone. In contrast, clone F3 showed a modest increase in H3K27ac enrichment upon CTCF loss consistent with the partial derepression of a subset of escapees in this clone [Extended Data Fig. 6d]. Accordingly, H3K27ac profiles across the six escapees within the *Mecp2–Hcfc1* cluster that showed increased Xi expression following CTCF depletion revealed a modest increase in H3K27ac enrichment at most promoter-associated and intergenic H3K27ac peaks [Extended Data Fig. 6d].

Altogether, these data demonstrate that, despite the presence of CTCF at active escapees, as well as at transition sites between active and inactive chromatin domains on the Xi, CTCF is dispensable for both the maintenance of escapee transcription and the preservation of these chromatin transitions in differentiated NPCs.

### Xi domain architecture is not driven by cohesin-mediated loop extrusion

We next investigated how the 3D TAD-like domains encompassing the *Mecp2-Hcfc1* and *Kdm5c* clusters on the Xi were affected by CTCF loss. To this end, we performed Capture Hi-C analysis of both clusters following 4 days of dTAG treatment in clones E6A7, F3 and C5C10. Overall, not all 3D structures on the Xi were lost, despite efficient depletion of CTCF as assessed by western blot and CTCF binding [Fig. 5a, Extended Data Fig. 7, Extended Data Fig. 8 a,b].

**Figure 5.**
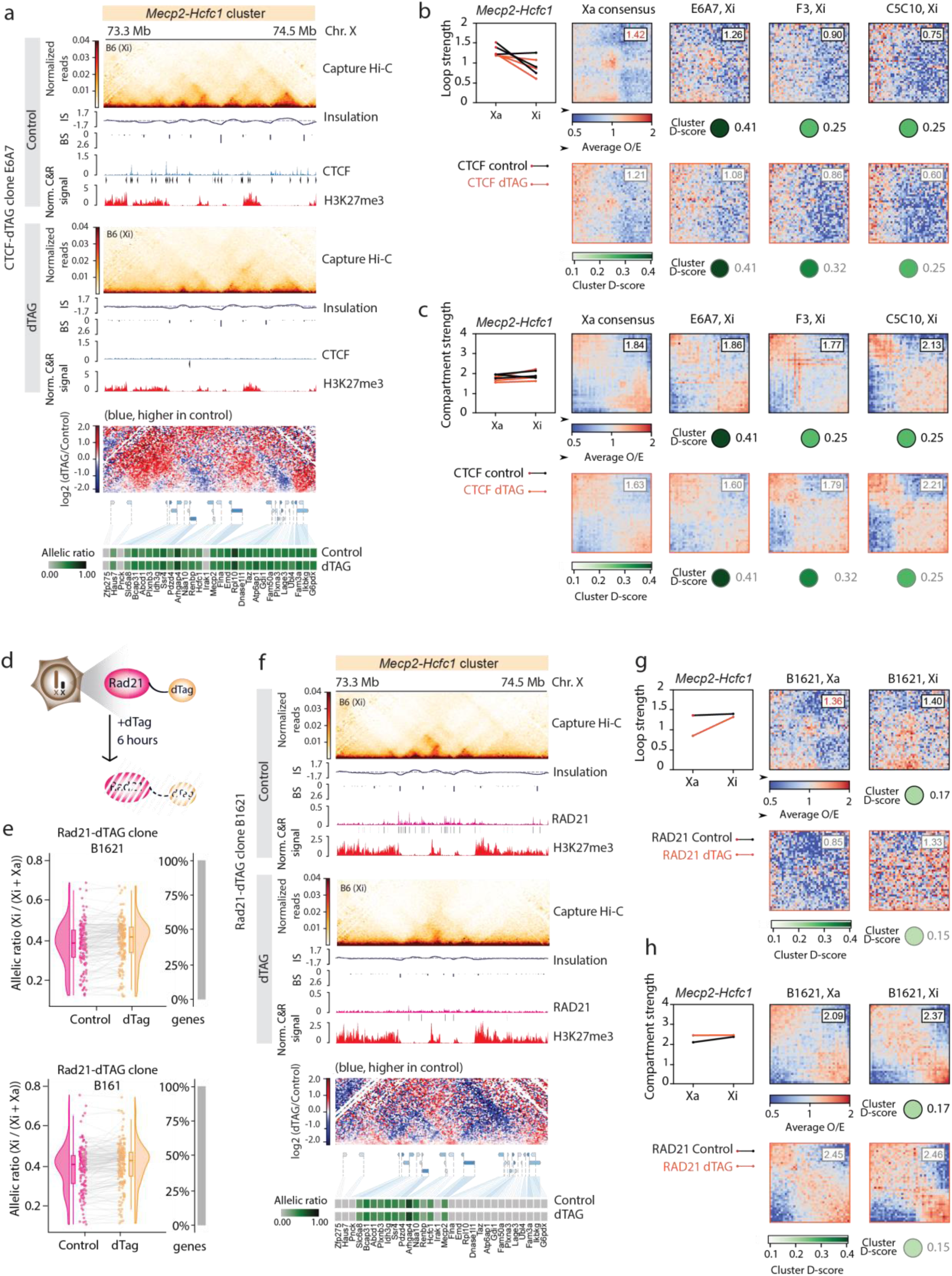
**a,** Capture Hi-C interaction maps, insulation scores, allele-specific CTCF CUT&RUN profiles and called peaks, and H3K27me3 CUT&RUN profiles across the *Mecp2–Hcfc1* escape cluster are shown for the CTCF-dTAG NPC clone E6A7 under control conditions and following dTAG treatment. Differential Capture Hi-C maps show changes in chromatin interactions between control and 4-day dTAG-treated samples. Capture Hi-C data are shown at 10 kb resolution. CTCF motif orientation is indicated by arrowheads. Heatmaps of allelic ratios for all genes within the cluster are shown for the control and dTAG-treated conditions. **b**, Loop strength at the *Mecp2*-*Hcfc1* cluster across CTCF-dTAG clones E6A7, F3 and C5C10, for the control (top) and dTAG-treated (bottom) conditions. Per-clone mean loop strength on Xa and Xi (top-left; one line per clone – control (black) and dTAG-treated (orange) conditions; loops were called on Xa), together with average observed-over-expected loop pile-up matrices at 5 kb resolution centered on consensus *Mecp2*-*Hcfc1* loops (see methods); mean O/E in the central 3 × 3 pixels is shown at top right of each pile-up. The Xa consensus pile-up uses the merged Xa cooler (see methods); Xi pile-ups are clone-specific (E6A7, F3, C5C10). Green circles below each pile-up reflect the per-clone average cluster D-score. **c**, As in **b** for compartment strength. Saddle plots are shown for the Xa consensus (see methods) and clone-specific Xi chromosomes. **d**, Schematic of the experimental design for acute RAD21 degradation in NPCs. **e**, Raincloud plots showing allelic ratios of X-linked genes with allelic ratio>0.1 in either the control or dTAG-treated condition across RAD21-dTAG NPC clones. Bar plots indicated the percentage of genes exhibiting a significant increase (green) or decrease (red) in allelic ratio following dTAG treatment (Δallelic ratio > 0.05; adjusted P < 0.05). No individual genes exhibited a significant change in allelic ratio across all clones. A small but significant increase in the mean escape level across all escapees was observed following dTAG treatment (mean ΔdTAG–control < 0.02; adjusted *P* ≤ 3.52 × 10⁻⁶). **f**, Capture Hi-C interaction maps, insulation scores, allele-specific RAD21 CUT&RUN profiles and called peaks, and H3K27me3 CUT&RUN profiles across the *Mecp2–Hcfc1* escape cluster are shown for the RAD21-dTAG NPC clone B1621 under control conditions and following dTAG treatment. Differential Capture Hi-C maps show changes in chromatin interactions between control and 6-hour dTAG-treated samples. Capture Hi-C data are shown at 10 kb resolution. Heatmaps of allelic ratios for all genes within the cluster are shown for the control and dTAG-treated conditions. **g**, As in **b** for RAD21-dTAG clone B1621. **h**, As in **g** for compartment strength. [IS - insulation score; BS - boundary score]

We quantified changes in loop strength upon CTCF loss across the *Mecp2-Hcfc1* and *Kdm5c* clusters on both the Xa and Xi, with the same strategy used to assess the 3D topology of these clusters across clones with different levels of escape [Fig. 5b,c, Extended Data Fig. 8c,d]. Specifically, we used the same consensus set of focal loops defined on the merged Xa Capture Hi-C maps as the reference coordinate system for pile-up analysis. First, we examined changes in loop strength upon CTCF depletion, on both the Xa and Xi in clones E6A7, F3 and C5C10 [Fig. 5b, Extended Data Fig. 8c]. This analysis revealed a consistent reduction in loop strength on both the Xa and the Xi at the *Mecp2-Hcfc1* and *Kdm5*c clusters following CTCF loss [Fig. 5b, Extended Data Fig. 8c], in agreement with previous reports that CTCF anchors cohesin-extruded loops genome-wide (Hsieh et al., 2022; Kubo et al., 2021; Nora et al., 2017).

We assessed compartment strength upon CTCF depletion at both the *Mecp2-Hcfc1* and *Kdm5c* clusters by performing saddle-plot analysis independently for the Xa and Xi before and after CTCF depletion with dTAG treatment [Fig. 5c, Extended Data Fig. 8d]. In contrast to loop strength, compartment strength remained essentially unchanged upon CTCF depletion at both clusters, and was even slightly reinforced on the Xi in clones C5C10 and F3 [Fig. 5c, Extended Data Fig. 8d].

These observations show that active and inactive domains at Xi escapee clusters remain segregated in the absence of CTCF, despite the loss of CTCF-anchored loops [Fig. 5 b-c, Extended Data Fig. 8c-d]. This is consistent with previous studies showing that A/B compartment organization occurs independently of CTCF (Nora et al., 2017; Rao et al., 2017; Schwarzer et al., 2017), and suggests that Xi 3D domains are likely driven by both loop extrusion and compartmentalization [Fig. 5 b-c, Extended Data Fig. 8c-d].

Since CTCF depletion primarily alters the positions at which chromatin loops are anchored, rather than cohesin-mediated loop formation itself (Davidson and Peters, 2021; de Wit and Nora, 2023; Rao et al., 2017) we additionally tagged the RAD21 cohesin subunit with the dTAG degron to definitively test whether 3D domains on the Xi depend on CTCF-cohesin-mediated loop extrusion [Fig. 5d; Extended Data Fig. 9a]. We integrated the FKBP12^F36V^ degron tag at the C terminus of RAD21 on both alleles of TX1072 ESCs by CRISPR-Cas9 targeting [Fig. 5d]. RAD21-dTAG ESCs were differentiated into NPCs and single cells were isolated and expanded to generate NPC clones B161 and B1621. We acutely depleted RAD21 for 6h, which resulted in robust depletion while avoiding adverse effects on NPC viability [Extended Data Fig. 9a-c] and performed RNA-seq analysis [Extended Data Fig. 9d]. Loss of RAD21 led to genome-wide effects at the transcriptomic level, with more genes being downregulated (368-411) than upregulated (172-216), consistent with observations from previous studies [Extended Data Fig. 9d] (Hsieh et al., 2022; Kriz et al., 2021; Rhodes et al., 2020).

The allelic ratios of escapees remained largely unchanged upon RAD21 depletion, with escapees remaining transcriptionally active on the Xi and silent genes not becoming reactivated [Fig. 5e]. In particular, although the general distribution of allelic ratios across escapees showed a modest increase in both clones, no individual gene reached statistical significance, likely because the effect size was too small to be detected at the single-gene level [Fig. 5e]. This is similar to the results obtained upon CTCF loss, which did not lead to silencing of escapees, but rather to an increase in their expression levels, at least in four of the clones analysed [Fig. 4]. Furthermore, allele-specific analysis of expression changes on the Xa and Xi upon RAD21 loss revealed the expression of only a minority of escapees affected, being either up- or downregulated. This occurred in most cases on both alleles rather than specifically on the Xi [Extended Data Fig. 9e]. We also checked Xist expression levels, which were unaffected by RAD21 depletion, again excluding any indirect effect on escapee expression [Extended Data Fig. 9f]. These results indicate that RAD21 is dispensable for escape maintenance in NPCs.

We then quantified changes in loop and compartment strength at the *Mecp2-Hcfc1* and *Kdm5c* clusters upon RAD21 depletion by Capture Hi-C [Fig. 5f, Extended Data Fig. 10]. Applying the same pile-up and saddle-plot framework to clone B1621, we observed attenuated loops at both the *Mecp2-Hcfc1* and *Kdm5c* clusters upon RAD21 depletion, on both the Xa and the Xi, in line with the established role of cohesin in loop extrusion [Fig. 5f, Extended Data Fig. 10] (Rao et al., 2017; Schwarzer et al., 2017). In contrast, RAD21 depletion further reinforced compartmentalization on the Xi at both the *Kdm5c* and *Mecp2-Hcfc1* clusters [Fig. 5h, Extended Data Fig. 10]. While compartment strength also increased on the Xa at the *Mecp2-Hcfc1* cluster [Fig. 5g], the effect at the *Kdm5c* cluster was largely restricted to the Xi, where the increase represented the greatest difference between control and dTAG-treated cells [Extended Data Fig. 10].

In summary, the 3D architecture of Xi escape domains is not driven by CTCF-cohesin loop extrusion, but likely reflects the compartmentalization of active and inactive domains on the Xi, independently of the action of both CTCF and RAD21.

### 3D domains on the Xi are spatially contiguous compartments of active chromatin

Given the above findings, we set out to explore the spatial chromatin organisation of escapee domains, in NPC clones showing different degrees of escape, by applying a METALoci spatial autocorrelation analysis (Mota-Gómez et al., 2026) to our allele-specific Capture Hi-C maps. Briefly, for each clone and allele, the top Hi-C contacts at the *Mecp2-Hcfc1* and *Kdm5c* loci were embedded in 2D using the Kamada-Kawai algorithm (Kamada and Kawai, 1989), placing frequently contacting bins in spatial proximity. CUT&RUN signals for H3K27ac and H3K27me3 were projected onto this layout, and local Moran’s I (Anselin, 1995) was computed at each bin to identify regions whose chromatin signal is significantly co-enriched with that of their 3D neighbours. Bins were classified into four quadrants: HH (high signal surrounded by high signal), LL (low surrounded by low), and the mixed LH and HL categories. Therefore, significant contiguous HH or LL bins mark spatial chromatin hubs either enriched or depleted of the particular chromatin mark. We use the combined fraction of significant HH and LL bins as a proxy for compartmentalisation strength and visualise the results as Gaudí plots, in which HH and LL regions appear as contiguous spatial domains [Fig. 6].

**Figure 6.**
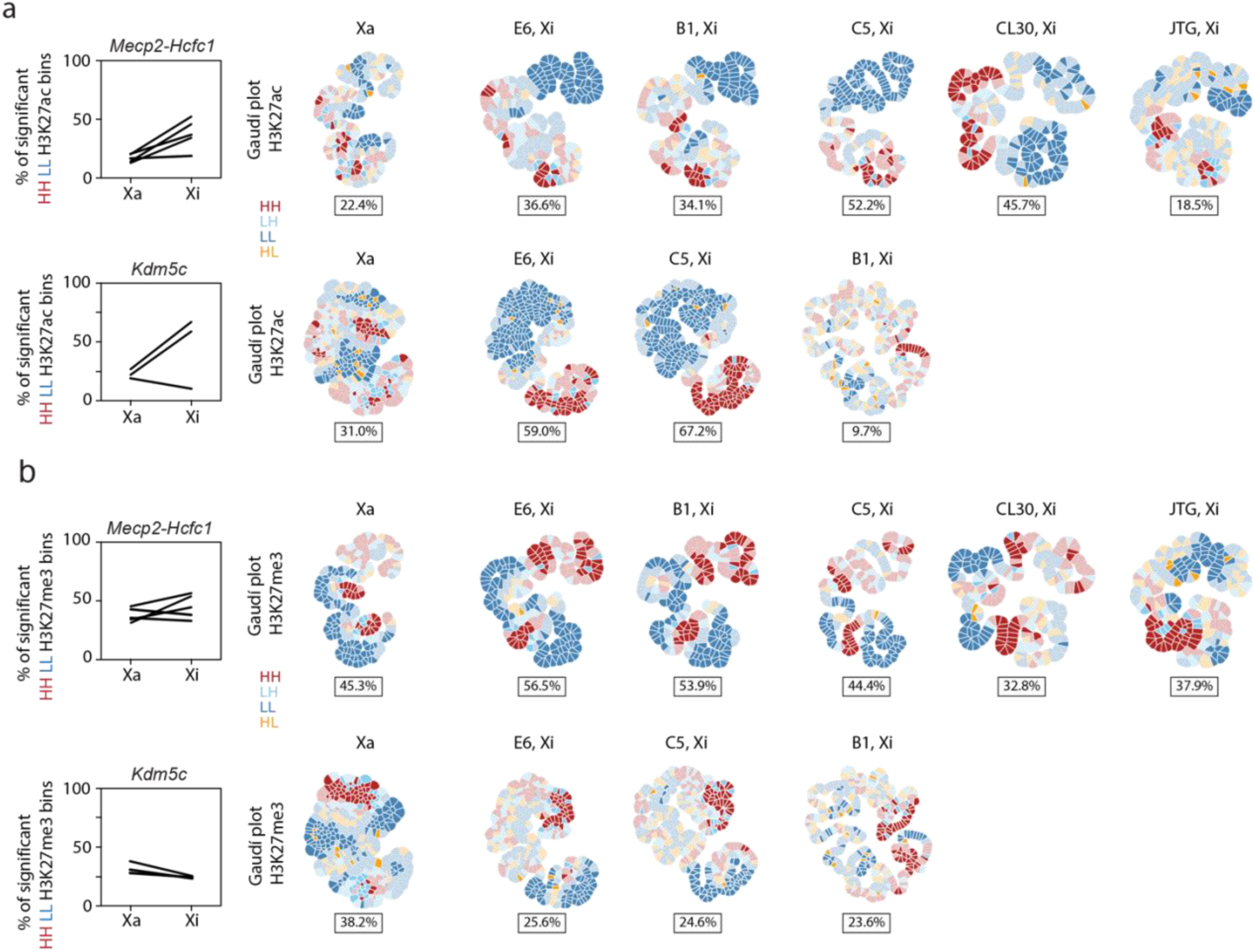
**a**, Gaudí plots (METALoci v1.3.2) for the *Mecp2*-*Hcfc1* and *Kdm5c* escapee cluster at 5 kb resolution, computed on the Xi in individual NPC clones and on the pooled Xa consensus (see methods). For each locus × clone × H3K27ac coverage, the Capture Hi-C contact map was embedded in 2D using the Kamada-Kawai algorithm (minimum interaction strength 1.5, persistence length 15) and signal-bin spatial autocorrelation was quantified by Local Moran’s I. Each polygon represents one 5 kb bin, coloured by its Moran’s quadrant: HH (red), LL (blue), HL (yellow) and LH (light blue). Subplot titles report the dataset and compartment-like strength, defined as the fraction of LMI-significant (p < 0.05) HH and LL bins: (N_HH,sig + N_LL,sig) / N_total × 100. Line plots (left) show the per-clone percentage of significant HH/LL H3K27ac bins on the Xa and Xi. **b**, As in **a** for H3K27me3.

On the Xa, H3K27ac signal shows low spatial autocorrelation for both *Kdm5c* and *Mecp2-Hcfc1* clusters across all clones [Fig. 6a]. This is consistent with a 3D architecture that is more loop extrusion-dominated for the Xa, where H3K27ac is present but spatially diffused and, therefore, does not clearly generate the local signal clustering detected by METALoci. In contrast, on the Xi, H3K27ac compartmentalisation increases markedly, and its magnitude tracks the degree of transcriptional escape at each locus.

At the *Kdm5c* cluster, this correspondence is not as clear for all clones. NPC clones E6 and C5, which retain substantial escape, display large compartmentalization of H3K27ac occupying 59% and 67% of significant bins on the Xi, respectively. Clone B1, in which only two genes escape from XCI [Fig. 1d], results in low compartmentalization of H3K27ac (9.7%). The *Mecp2-Hcfc1* cluster follows the same logic. H3K27ac hub formation is most extensive in clones C5 and CL30 (52% and 46%), whereas in clone JTG (in which only three genes escape at this locus [Fig. 1c]) the fraction of significant bins is low (18.5%) and does not increase relative to the Xa, in line with the weak compartmentalisation strength independently detected by Hi-C saddle analysis in this clone [Fig. 3b].

H3K27me3 shows a very complementary pattern to H3K27ac [Fig. 6b], occupying the chromatin territory spatially excluded by the H3K27ac hub in each clone. The two marks are mutually exclusive within the locus, and the boundary between their respective 3D hubs coincides with the limits of the escape domain as defined independently by allele-specific RNA-seq [Fig. 1c–d] and H3K27me3 valley calling [Fig. 1e, Extended Data Fig. 2c,f,].

Together, these results show that the 3D domains characterising escapee clusters on the Xi, are spatially contiguous compartments of active chromatin, maintained by the self-organisation of H3K27ac-enriched bins in 3D, rather than by CTCF- or cohesin-anchored loop structures. Furthermore, the boundaries of these compartments correspond to the segregation between active and Polycomb-silenced chromatin.

## Discussion

In this study, we have investigated what underlies domains of facultative escape on the Xi by exploring their 3D organisation, transcriptional and chromatin states in clonal, genetically identical NPC lines with varying levels of escape. This has revealed that variable facultative escape is accompanied by changes in 3D topology and chromatin state, with a tight coupling between these regulatory layers. Our study has also addressed several long-standing questions regarding the role of CTCF in promoting escape from XCI, including whether CTCF promotes escape through its architectural functions, acts as an insulator to prevent the spread of XCI into escape domains, or directly contributes to escapee genes transcription (Berletch et al., 2015; Filippova et al., 2005; Giorgetti et al., 2016; Goto and Kimura, 2009). The generation of CTCF and RAD21 degron lines, using genetically identical NPC clones with different levels of facultative escape, allowed us to assess the effects of disrupting loop extrusion within escapee domains on the Xi, as well as the consequences of this on gene expression within the domain and at neighbouring regions on the Xi.

We found that near-complete depletion of either CTCF or RAD21 did not result in a complete loss of escape-domain architecture. Instead, disruption of loop extrusion primarily attenuated looping strength, while compartments were maintained following CTCF depletion and, in the case of RAD21 depletion, often became more pronounced. This is in line with recent work proposing that enhancer-promoter interactions can be driven by compartmentalization mechanisms and that, in fact, the microcompartments underlying these interactions are largely unaffected by the loss of cohesin-mediated loop extrusion (Goel et al., 2023). Importantly, it had previously been proposed that TAD-like structures encompassing domains of escape on the Xi, which is otherwise globally depleted for TADs, might be the result of cohesin-mediated loop extrusion (Giorgetti et al., 2016). The present analysis shows that 3D escapee domains are not defined by a single architectural feature. Rather, these domains result from an interplay between looping interactions and compartmentalization, supporting the emerging view that these are distinct but coexisting organizational principles, rather than different levels of a hierarchical folding architecture (Goel et al., 2023; Harris et al., 2023). The fact that 3D escapee domains remain detectable following depletion of either CTCF or RAD21, despite the attenuation of looping interactions, indicates that these domains represent fine-scale, activity-associated chromatin compartments rather than loop-extruded domains. Consistent with this finding, recent studies have shown that compartmentalization can occur at remarkably fine scales, overlapping with loop-sized domains and even distinguishing the 5′ and 3′ ends of genes (Harris et al., 2023).

Models for escape from XCI have evoked multiple potential roles for CTCF: as an insulator that enables genes to remain expressed on the otherwise silent X chromosome; as a facilitator of escape through looping out of the Barr body; as a barrier to spreading of escape or silencing (Berletch et al., 2015; Ciavatta et al., 2006; Fang et al., 2025; Filippova et al., 2005; Giorgetti et al., 2016; Goto and Kimura, 2009). Remarkably transcription of escapees from the Xi remained largely unaffected following depletion of either CTCF and RAD21, independently of the number of escapees and their expression levels. If anything, loss of CTCF was associated with a modest upregulation of escapees on the Xi in clones with lower escape levels. Moreover, we found no evidence for reactivation of silenced genes or disruption of the transitions between active and inactive chromatin that define putative escape-domain boundaries on the Xi. These transition regions, characterized by enrichment of H3K27ac within active domains and its sharp decline across adjacent inactive regions enriched for H3K27me3, remained intact despite the presence of CTCF at many of these sites.

Thus, although CTCF and cohesin contribute to the structural organization of escape domains, neither CTCF nor cohesin-mediated loop extrusion are required for the maintenance of escape transcription or insulation of escape domains in NPCs. Interestingly, previous work reported reinforcement of long-range “superloops” between escapee loci across the Xi following RAD21 degradation, including interactions involving *Kdm5c*, *Ogt*, *Bcor* and *Utp14a* (Kriz et al., 2021). Because these interactions were strengthened rather than lost upon RAD21 depletion, they were proposed to reflect chromatin interactions established through mechanisms other than cohesin-mediated loop extrusion (Kriz et al., 2021), which is consistent with our findings. Furthermore, the interactions of active TAD-like domains encompassing escapees along the Xi was also observed in extraembryonic tissues (Du et al., 2024), again supporting the notion that they might represent active chromatin compartments.

Our METALoci analysis shows that these escapee interactions on the Xi are indeed most likely due to compartmentalization of active chromatin, with escape domains scaling with escape activity independently of their genomic context on the Xi. Importantly, this is a feature of escapee domains exclusively on the Xi, not on the Xa for the same regions. We further show that escapee domains are driven by the spatial clustering of H3K27ac-enriched chromatin; that they persist or are reinforced following disruption of loop extrusion; and that their boundaries are defined by transitions between opposing chromatin states. The association we found between H3K27ac-enriched chromatin and escape-domain compartmentalization underlies the broader relationship between chromatin state and 3D genome topology. Indeed, histone modifications are among the strongest predictors of compartment identity and chromatin interactions in computational models of genome folding (Feng et al., n.d.; Harris and Rowley, 2024; Piecyk et al., 2022). Moreover, hyperacetylated chromatin has been shown to engage in long-range interactions that form compartment-like structures, suggesting that active chromatin states can contribute to spatial chromatin segregation (Rosencrance et al., 2020). While these observations are consistent with a role for H3K27ac-enriched chromatin in driving escape-domain compartmentalization, whether histone modifications actively drive compartment formation or instead reflect underlying transcriptional activity in these domains remains unresolved. Our recent study exploring the impact of increasing Xist RNA levels on escape in NPCs, demonstrated that these 3D compartments are in fact lost upon Xist RNA increase (Hauth et al., 2026). However this effect is seen only in the presence of SPEN, which is a key partner of Xist and regulator of XCI (Dossin et al., 2020). Given that SPEN, via its SPOC domain, recruits chromatin modifiers including H3K27 deacetylase HDAC3 (Dossin et al., 2020), we propose that 3D escape compartments on the Xi are indeed H3K27ac chromatin-dependent. Furthermore, as highlighted by Dekker and Mirny (Dekker and Mirny, 2024), histones may play a fundamental role in compartment formation. Chromatin compartmentalization is readily detected across eukaryotes with canonical nucleosome-based chromatin organization, whereas it appears to be absent from systems lacking canonical nucleosomes, such as dinoflagellates and some archaea (Dekker and Mirny, 2024). Consistent with this view, recent comparative analyses across holozoans indicate that compartmentalization and focal chromatin looping are evolutionarily and mechanistically separable features of genome organization (Kim et al., 2025).

In conclusion, we have shown that neither CTCF and RAD21 have a major role in maintaining escapee status of genes on the Xi. Our study demonstrates that compartmentalization of active chromatin on the heterochromatic Xi, rather than loop extrusion, is the major organizational feature underlying escape domains on the Xi. This work also highlights the remarkable stability of active chromatin compartments encompassing escapee genes, on an otherwise inactive chromosome. An open question concerns the way in which these domains are initially established following XCI, and what defines their frontiers. This remains an exciting area of future study.

## Methods

### Cell culture

The TX1072 (*Mus musculus castaneus* (Cast/EiJ) × *Mus musculus domesticus* (C57BL/6J)) mESC line was previously derived in the Heard lab (Schulz et al., 2014). Cells were cultured in 2i-containing ES cell media (DMEM, 15% fetal bovine serum, 0.1 mM β-mercaptoethanol, 1,000 U mL^−1^ leukaemia inhibitory factor, CHIR99021i (3 µM) and PD0325901 (1 µM)) on gelatine-coated (0.1% gelatine in 1× PBS) cell culture dishes.

### NPC differentiation

TX1072 mESCs carrying a doxycycline (Dox)-inducible promoter upstream of the *Xist* endogenous locus on the C57BL/6 X chromosome were differentiated and sub-cloned as previously described (Gendrel et al., 2014; Hauth et al., 2026). In brief, 1 × 10^6^ ES cells were seeded in a gelatin-coated 10-cm petri dish in N2B27 media (DMEM/F12:Neurobasal (1:1), L-glutamine, 0.1 mM 2-mercaptoethanol). At day 7 of differentiation, 3 × 10^6^ cells were plated in N2B27 media supplemented with epidermal growth factor (EGF) and fibroblast growth factor (FGF) (10 ng ml^−1^ each) in bacterial Petri dishes to prevent cells from attaching to the plates. After 3 days of cell growth in suspension as cellular aggregates (or spheres), the aggregates were harvested and plated onto gelatine-coated 10-cm Petri dishes in N2B27 media supplemented with EGF and FGF. Monolayer NPCs grew out of the attached sphere. To generate NPC clones E6, B1, C5, JTG, CL30 and the RAD21-dTAG clones, 5,000–10,000 single cells were plated in 10-cm Petri dishes and single clones were manually picked after 15–20 days. Single NPC clones were expanded and characterized by RNA-FISH for *Xist* and *Huwe1* to assess karyotype stability. In case of unstable karyotype resulting in gain or loss of chromosomes, NPC clones were either discarded or further sub-cloned. Clones E6, C5 and B1 were derived upon ESCs differentiation in absence of Dox. Clones CL30, JTG, and all CTCF- and RAD21-degron clones, were differentiated in the presence of Dox for the first 3-7 days of ESCs differentiation. Clone CL30 carries endogenous SPEN alleles tagged with AID– Halo and was previously generated in the laboratory (Dossin et al., 2020). NPCs were cultured on a gelatine-coated (0.1% gelatine in 1× PBS) cell culture dish in NPC media (N2B27 media supplemented with with FGF2 (10 ng/mL) and EGF (10 ng/mL); both from PeproTech). All NPC clones were regularly tested for karyotype stability by RNA-FISH at the end point of each experiment, before sequencing.

### Cell treatments

NPCs were treated with dTAG-13 (500 nM in culture media, *SML2601-1MG, Sigma*) one day after plating to achieve acute protein depletion; dTAG-containing media was changed every 24-48 h.

### Plasmid construction

Single-guide RNAs (sgRNAs) targeting the endogenous *Ctcf* and *Rad21* loci were cloned into the pX330 vector (Addgene plasmid #42230), which encodes *Streptococcus pyogenes* Cas9, by BbsI-mediated digestion followed by ligation of annealed oligonucleotides. The sgRNA target sequences were as follows: *Ctcf*, 5′-ATGATGGACCGGTGATGCTG-3′; *Rad21*, 5′-CCACGGTTCCATATTATCTG-3′ (pX330-EN1082_Rad21_STOP, Addgene #156450). For CTCF tagging, a targeting construct (pTarget-CTCF-EGFP-dTag-HygR) was generated by inserting the sequence coding for EGFP, short peptide linker sequences and FKBP12^F36V^ (dTAG) degradation domain tags in frame with the *Ctcf* coding sequence followed by the *Ctcf* stop codon as well as a hygromycin resistance cassette flanked by FRT sites. The design of the homology arms was based on a previously described CTCF-AID-eGFP targeting vector (pEN84; Nora et al., 2017)(gift from the Nora lab). For NPC targeting, the HygR selection cassette was removed from the donor construct by InFusion cloning. To tag the C terminus of endogenous RAD21 with the FKBP12^F36V^ (dTAG) degradation domain, a previously described targeting vector was used (Rad21-HaloTag-FKBP, (Mach et al., 2022)(gift from the Giorgetti lab).

### Genomic engineering of mESCs and NPCs

CRISPR–Cas9-mediated genome editing in mESCs and NPCs was performed using nucleofection (4D-Nucleofector, Lonza). Briefly, 5 million cells were nucleofected with 10 µg total DNA comprising targeting vectors and guide RNA-Cas9 encoding plasmids. ESCs were plated at limiting dilution and selected with hygromycin (250 µg/mL) starting 24-48 h post-transfection. For flippase-mediated removal of resistance cassettes no selection was applied. Individual ESCs colonies were picked after 5–7 days and expanded for PCR genotyping. NPCs nucleofected with plasmid pTarget-CTCF-EGFP-dTag-HygR were single-cell sorted 24-48 h post-transfection. Single clones were expanded for PCR genotyping. Karyotype stability was also assessed by RNA FISH.

### Protein extraction and western blotting

Total protein was extracted from >1 million cells. Cell pellets were washed in PBS, frozen at -80 °C, and lysed in RIPA buffer (50 mM Tris-HCl pH 8.0–8.5, 150 mM NaCl, 1% Triton X-100, 0.5% sodium deoxycholate, 0.1% SDS) supplemented with protease inhibitors (Roche). Lysates were incubated on ice for 1 h, briefly sonicated (Bioruptor, Diagenode), and cleared by centrifugation at 4 °C (40 min, > 10000g). Protein concentration was determined using a Bradford assay (Bio-Rad), and samples were normalized and denatured in LDS sample buffer (Invitrogen) at 95 °C for 5 min. Equal amounts of protein (10-20 µg) were separated on 4-12% Bis-Tris gels (Invitrogen) and transferred to nitrocellulose membranes by wet transfer in Tris-glycine buffer containing 10% methanol. Membranes were stained with Ponceau S to confirm transfer, blocked in 5% milk in PBS containing 0.3% NP-40, and incubated overnight at 4 °C with primary antibodies: CTCF (61311, Active Motif, 1:1000), Rad21 (ab154769, Abcam, 1:1,000) and TBP (ab300656, Abcam, 1:1000). After washing, membranes were incubated with HRP-conjugated secondary antibodies (Cytiva; 1:5000) for 1 h at room temperature. Signal was detected using LumiLight WB substrate (Roche) and imaged on a ChemiDoc system (Bio-Rad).

### Flow cytometry and cell sorting

NPCs were harvested by dissociation into single-cell suspensions using accutase, collected by centrifugation and resuspended in cell sorting buffer (1× PBS supplemented with 2.5 mM EDTA (pH 8.0) and 2% (w/v) BSA (Merck, A4503-100G)). Cell suspensions were filtered through a 40-µm cell strainer to remove aggregates, stained with DRAQ7 viability dye (BioStatus, DR77524), and maintained on ice prior to flow cytometric acquisition. Flow cytometric analysis was performed using a BD LSRFortessa™ flow cytometer (BD Biosciences). For single-cell sorting of NPCs, individual cells were sorted into matrigel-coated 96-well plates (Corning, 356234). DRAQ7 was used for live cell labelling and GFP fluorescence measured successfully targeted cells (CTCF-GFP-dTAG). DRAQ7 fluorescence was detected using the 640 nm excitation laser and a 670/14 nm emission filter, whereas GFP fluorescence was detected using the 488 nm excitation laser and a 530/30 nm emission filter. For measuring the degradation of CTCF in the CTCF-GFP-dTAG lines, instrument settings were maintained constant across all samples within an experiment. Flow cytometry data were processed and analyzed using FlowJo (BD Biosciences), and data were subsequently imported into R for downstream analysis and plot generation. Cells were initially gated based on forward scatter area (FSC-A) and side scatter area (SSC-A) to exclude debris. Doublets were excluded using SSC-A versus SSC-H gating, and dead cells were removed based on DRAQ7 fluorescence. GFP-positive cells were identified based on fluorescence intensity relative to GFP-negative controls, including dTAG-13-treated CTCF-degron NPCs where applicable. Identical gating thresholds were applied across all samples. The mean GFP fluorescence intensity was quantified in arbitrary units (a.u.) within the live cell population.

### RNA sequencing

RNA was extracted from >1 million NPCs using the RNeasy kit with on-column DNAse digestion (QIAGEN). RNA integrity was assessed using the Bioanalyzer Nano Kit and only high-quality RNA was used for library preparation. RNA was isolated using the NEBNext Poly(A) mRNA Magnetic Isolation Module (E7490L, New England Biolabs), and strand-specific libraries were generated using the NEBNext Ultra II Directional RNA Library Prep Kit (E7760L, New England Biolabs) on a Beckman i7 liquid handling platform. Libraries that passed the quality control were pooled in equimolar amounts and sequenced on NextSeq500 or 2000 (Illumina; 50-75 bp paired-end, >55 million reads/library). All RNA-seq experiments were performed in two independent biological replicates.

### Capture Hi-C

Capture Hi-C was performed as previously described (Hauth et al., 2022, 2026). Previously described biotinylated RNA probe arrays were used to tile two 3 Mb target regions on the mouse X chromosome: *Hcfc1*–*Mecp2* cluster (ChrX: 72,590,000–75,430,000) and *Kdm5c* cluster (ChrX: 150,210,000– 153,045,000) (Hauth et al., 2026). Capture Hi-C maps are from a single replicate, except for CTCF-degron clone F3, for which two independent biological replicates were performed per condition (Ctrl and dTAG).

### CUT&RUN

CUT&RUN experiments were performed as previously described (Hauth et al., 2026), with the exception of library preparation. In brief, 0.5 × 10^6^ cells were collected, permeabilized, and immobilized on Concanavalin A-coated magnetic beads. Nuclei were incubated overnight at 4 °C with primary antibodies (1:100). The following antibodies were used: H3K27me3 (9733, Cell Signaling), H3K27ac (39685, Active Motif), CTCF (61311, Active Motif) and RAD21 (ab154769, Abcam). After washing, samples were incubated with pA-MNase (generated by EMBL PepCore facility) at 700 ng/mL for 1 h at 4 °C. Chromatin digestion was initiated by addition of CaCl_2_ and performed for 30 min on ice. The reaction was quenched and following RNAse A and Proteinase K treatment, DNA was extracted by phenol-chloroform purification, ethanol precipitated, and size-selected using Ampure XP beads. Libraries were prepared from up to 250 ng DNA using the AcceI-NGS2S Plus DNA library kit (Swift Bioscience, 21024-SWI). For samples with low expected DNA yield (e.g. CTCF or Rad21 CUT&RUN in degron conditions), all available input was used for library preparation. Wild-type and CTCF-degron NPC clone F3 were sequenced at 50-75 bp paired-end on NextSeq500 or 2000. CTCF-degron clone E6A7 and RAD21-degron clone B1621 were sequenced at 75 bp paired-end on Element Biosciences AVITI System. All CUT&RUN experiments were performed in two independent biological replicates.

### Bioinformatics

#### Allele-specific RNA-seq data pre-processing

RNA-seq data were processed as previously described (Hauth et al., 2026) using a Nextflow pipeline developed for this project and available at https://github.com/yuviaapr/allele-specific_RNA-seq. Briefly, adapter trimming was performed using Trim Galore (https://github.com/FelixKrueger/TrimGalore) (version 0.6.3). For allele-specific mapping, reference genomes were generated by masking known C57BL/6 and CAST/EiJ heterozygous variants (https://ftp.ebi.ac.uk/pub/databases/mousegenomes/REL-2112-v8-SNPs_Indels/mgp_REL2021_snps.vcf.gz) with ambiguous bases (N) using bedtools maskfasta (Quinlan and Hall, 2010) (version 2.29.2), followed by genome indexing with STAR (Dobin et al. 2013) (v2.5.3a). Reads were aligned to the N-masked genomes using STAR with the following parameters: -- sjdbOverhang 99, --outFilterMultimapNmax 1, –outFilterMismatchNmax 999–, -- outFilterMismatchNoverLmax 0.06, --alignIntronMax 500000, --alignMatesGapMax 500000, and -- alignEndsType EndToEnd. All samples were aligned to the mm10 (GRCm38) reference genome. Reads mapping to the mitochondrial genome were excluded, and allele-specific assignment was performed using SNPsplit (Krueger and Andrews, 2016) (version 0.3.4) and the heterozygous variants. Gene-level read counts, both allele-specific and total, were generated using featureCounts(Liao et al., 2014) (v2.0.1).

#### Computation of allelic ratios in RNA-seq for escapee definition

Allele-specific expression per gene was calculated from allele-specific read counts. Lowly expressed genes (average allelic read count sum <10) were excluded. Escape from X-chromosome inactivation was defined based on the fraction of reads derived from the inactive X chromosome (Xi / (Xi + Xa), allelic ratio). Genes were classified as escapees if the allelic ratio exceeded 0.1 in both biological replicates. Xist was excluded from this analysis due to its role in X-chromosome inactivation. Genes with allelic ratios >0.8 were excluded from downstream analyses, as previously described (Hauth et al., 2026). Compared to our previous study, a small number of additional genes were classified as escapees in the present analysis. This difference reflects the lower number of biological replicates and the requirement that the allelic ratio threshold (> 0.1) be met in each replicate individually. Genes uniquely identified as escapees in this dataset were generally characterized by low overall expression levels, allelic ratios close to the defined threshold, or technical limitations such as reduced mappability or potential mapping biases at specific SNP positions.

#### Differential expression analysis

Differential expression analyses were performed using DESeq2 (Love et al., 2014)v1.34.0). Gene-level raw read counts were used as input, and lowly expressed genes (total counts <10 across NPC clone samples) were excluded. For total expression analyses, size factors were calculated separately for each clone excluding X-linked genes, and genes in chromosomes 1, 8 and 16 that tended to show aneuploidies in NPC clones. Dispersions were estimated using default methods, and differential expression was assessed using Wald tests between control and dTAG-treated samples. Genes with an absolute log₂ fold change (dTAG/control) > 1 and an adjusted *P* value < 0.05 were considered differentially expressed. Results from pairwise comparisons were extracted using the lfcShrink function with the *apeglm* estimator. For allele-specific analyses, read counts were separated by allele and analyzed using the same approach. Size factors estimated from total counts were applied to allele-specific datasets. Significance of allelic ratio differences between control and dTAG samples were tested using DESeq2 on raw allelic read counts, with a design to consider variation across condition, replicates and alleles (e.g. ∼condition + condition:replicate + condition:allele). Size factors were set to 1, as total coverage per gene normalizes allelic ratios within samples, eliminating library size effects. Wald test results were extracted using the *results* function with IHW p-value adjustment(Ignatiadis et al., 2016). As for the previous analyses, genes with <10 total counts across samples were filtered a priori, and results of genes with raw read counts in the lowest 0.15 percentile in at least one of the biological replicates were discarded.

#### Allele-specific Capture Hi-C analysis

Allele-specific Capture Hi-C data were processed using the HiC-Pro pipeline(Servant et al., 2015) (v2.11.4) as previously described(Hauth et al., 2026). Briefly, paired-end reads were aligned to an N-masked reference genome using Bowtie2(Langmead and Salzberg, 2012) and assigned to C57BL/6J or Cast/EiJ alleles. Low-quality reads, multi-mapping reads, singletons, and read pairs that did not map to the same genotype were then removed. Valid interaction pairs within the captured regions were retained and used to generate contact maps at 5 and 10 kb resolution using the cooler package(Abdennur and Mirny, 2020) (v0.8.9). Insulation scores were calculated using cooltools (https://github.com/open2c/cooltools, v0.3.2). Data visualization was performed using pyGenomeTracks(Lopez-Delisle, 2025). Summary statistics are provided in Supplementary Table 1.

#### Allele-specific CUT&RUN analysis

CUT&RUN data were processed as previously described (Hauth et al., 2026)using a Nextflow pipeline developed for this project (https://github.com/yuviaapr/allele-specific_CUTandRUN). Paired-end reads were aligned to an N-masked reference genome (see Allele-specific RNA-seq data pre-processing) using Bowtie2 (v2.3.4.1) with the parameters --very-sensitive, -X 1000, --no-mixed and --no-discordant. Low-quality and mitochondrial reads were removed using Samtools (Li et al., 2009)(v1.9). Allele-specific assignment was performed using SNPsplit (v0.3.4), followed by duplicate removal with Picard Tools (“Picard Toolkit.” 2019. Broad Institute, GitHub Repository. https://broadinstitute.github.io/picard/; Broad Institute) (v2.20.8). Separate BAM files corresponding to C57BL/6J, Cast/EiJ and total reads (allelic reads plus unassigned) were generated for downstream analyses. CUT&RUN libraries were normalised separately for each target (H3K27me3, H3K27ac, CTCF, Rad21) to render signal comparable across clones and genotypes. One replicate of CUT&RUN in CL30 targeting H3K27ac (CL30 rep2) was excluded from the normalisation, due to its poor correlation with the corresponding CL30 rep1 for the same mark. The total reads (“Gall”) BAM file together with its two corresponding allele-specific BAM files (G1, G2) were treated as a triplet sharing a single normalisation factor. Normalisation factors were derived with csaw v1.42.0(Lun and Smyth, 2016) on the total (“Gall”) BAM files of all clones for a given mark, run under R v4.5.1. BAMs were read with csaw::readParam configured for paired-end data (pe = “both”), with a maximum fragment size of 300 bp and the ENCODE mm10 blacklist v2(Amemiya et al., 2019) excluded via the discard argument. Read counts were restricted to autosomes (chr2-5, chr7-10, chr13-15, chr17-19) and summarised in non-overlapping 10 kb bins with csaw::windowCounts (bin = TRUE, width = 10000).

Per-sample normalisation factors were then computed with csaw::normFactors using the trimmed mean of M-values (TMM) method(Robinson and Oshlack, 2010) from edgeR v4.6.3(Robinson et al., 2010). BAM I/O and blacklist handling used Rsamtools v2.24.0(Morgan et al., n.d.) and rtracklayer v1.68.0(Lawrence et al., 2009). The normalization factor for each replicate was then propagated to the two corresponding allelic BAM files. Normalised bigWig tracks were generated with deepTools bamCoverage v3.5.5(Ramírez, 2016) using --binSize 10, --normalizeUsing None, and an explicit scale factor -- scaleFactor = 1e6 / (LibSize × NormFactor), combining CPM scaling with the csaw/edgeR-TMM correction (LibSize, mapped-read count of the total BAM; NormFactor, the csaw normFactors output). Reads were extended to their fragment length with --extendReads; the point-source factors CTCF and Rad21 were additionally recentred with --centerReads. Allele-specific bigWigs (G1, G2) were generated using the same scale factor as their parent total BAM, following the convention established previously(Żylicz et al., 2019). Replicate bigWigs were merged per clone with deepTools bigwigCompare v3.5.2(Ramírez, 2016) (--operation mean at 20 bp and 5 kb bin sizes). For peak calling in CTCF, Rad21 and H3K27ac samples sequencing depth was normalized by subsampling reads within a twofold range of the lowest library size prior to alignment using seqtk (https://github.com/lh3/seqtk) (v1.3; -s123) and peaks were called from the total reads bam file (allelic and unassigned reads) for each replicate separately using macs2(Zhang et al., 2008) (v2.2.7.1) without the use of a control or mock data file. Consensus peak sets for each biological replicate pair were generated by retaining peaks overlapping by at least 1 bp using bedtools (v2.29.2). A union peak set for NPC samples was constructed by merging overlapping regions from clone-specific consensus peak files using bedtools. Peaks were annotated to genomic features using ChIPseeker (v1.34.1)(Wang et al., 2022) and grouped into three categories: promoter, genebody and intergenic. Eac peak was additionally assigned to the nearest gene. Motif analysis of CTCF peaks was performed using the JASPAR2020 database(Fornes et al., 2020) and only CTCFp peaks overlapping a motif were kept for downstream analysis. Read counts within union peak regions were quantified from allele-specific BAM files using featureCounts(Liao et al., 2014) (v2.0.1), and allelic ratios were calculated as Xi / (Xi + Xa). Downstream analysis of allelic ratios considered only peaks that were originally called in the respective clone. CTCF peaks were classified as biallelic if their allelic ratio exceeded 0.25 and as Xa-monoallelic otherwise.

#### Loop calling

Chromatin loops were called with Chromosight v1.6.3(Matthey-Doret et al., 2020) on two Xa merged .mcool files (Kdm5c_Xa and Mecp2_Xa), each obtained by pooling all NPC samples used in this study. For the *Mecp2-Hcfc1* cluster: 9 clones in total (WT clones E6, B1, C5, JTG, CL30, CTCF-dTAG clones E6A7, F3, C5C10 in untreated conditions, RAD21-dTAG clone B1621 in untreated conditions). *Kdm5c* cluster: 6 clones in total (WT clones E6, B1, C5, CTCF-dTAG clones E6A7, C5C10 in untreated conditions, RAD21-dTAG clone B1621 in untreated conditions) at 5 kb resolution. The “loops” pattern was used with--norm auto, --min-dist 50,000 bp, --max-dist 5,000,000 bp, --pearson 0.3, --perc-undetected 40, --perc-zero 10, --n-mads 3 and --min-separation 10,000 bp (2 × resolution). Residual false-positive calls were removed manually from each loop set after visualization. Loop strength at individual loops was defined as the mean observed-over-expected contact frequency in the 3 × 3 bin square centred on each loop anchor, with expected values derived from the genome-wide distance-decay profile (cooltools.expected_cis). For each clone, per-loop strength was computed against the corresponding individual Xa or Xi cooler and then averaged to yield a single mean loop strength per clone. Violin plots show the distribution of these per-clone mean values across clone groups; pile-up matrices were generated with coolpuppy v1.1.0(Flyamer et al., 2020) by stacking observed-over-expected sub-matrices across all consensus loop positions; the reported pile-up score is the mean O/E in the central 3 × 3 pixels.

#### Compartment calling

Chromatin compartments were called with cooltools v0.5.4(Open2C et al., 2024) on individual and merged Xa (for WT and CTCF-dTAG clones in untreated condition for both clusters the same pooling strategy was used as for loop calling (see above); for CTCF-dTAG clones in treated condition clones E6A7, F3 and C5C10 were pooled for the *Mecp2-Hcfc1* cluster and clones E6A7 and C5C10 were pooled for the *Kdm5c* cluster; for RAD21-dTAG clone B1621 the individual .mcool files were used) and individual Xi ICE-balanced .mcool files at 5 kb resolution, using cooltools.eigs_cis with the mm10 GC-content track (computed with bioframe v0.8.0, bioframe.frac_gc) as the phasing reference. The first three eigenvectors (E1-E3) were visually inspected per cooler, and the one giving the clearest checkerboard pattern was retained. Saddle plots were computed with cooltools.saddle on cis observed/expected matrices (cooltools.expected_cis), using N = 38 equally spaced eigenvalue bins between the 2.5th and 97.5th percentiles (ranging from 0.025 to 0.975). Compartment strength was quantified as (AA + BB) / (AB + BA), taking the 8 most extreme bins per side (top-left and bottom-right for AA and BB; top-right and bottom-left for AB and BA), equivalent to cooltools.api.saddle.saddle_strength at extent = 8.

#### METALoci analysis

Gaudí plots were generated with METALoci v1.3.2(Mota-Gómez et al., 2026) using merged bigWig tracks as signal input and corresponding .mcool files as Hi-C input, at 5 kb resolution. Two complementary inputs were used: (i) per-clone files, in which only replicates were merged; and (ii) consensus Xa and Xi files, in which both replicates and clones were merged. Signal tracks comprised H3K27ac and H3K27me3. The METALoci pipeline was run with default parameters unless indicated. First, *metaloci prep* was used to parse the Hi-C and CUT&RUN datasets at 5kb resolutions. Second, *metaloci layout* was run with default parameters at 5Kb resolution with the parameters “-o” set at 1.5 and “-pl” set at 15. Third, *metaloci lm* was run with default parameters to calculate the local Moran’s I of each input signal parsed in the first step. And fourth, *metaloci figure* with default parameters was run to generate Gaudí plots and summary figures for each signal. Compartment-like strength reported from Gaudí plots was calculated as the fraction of LMI-significant HH and LL bins in the locus: compartment-like strength (%) = (N_HH,sig + N_LL,sig) / N_total × 100, where N_HH,sig (N_LL,sig) is the number of bins in Moran’s quadrant 1 and 3 with LMI p-value < 0.05, and N_total is the total number of bins in the analyzed genomic region.

#### Identification of H3K27me3 valleys on the inactive X chromosome

H3K27me3 valleys on the Xi were called independently for each clone from the per-clone, replicate-averaged H3K27me3 bigWig tracks (deepTools(Ramírez, 2016) bigwigCompare v3.5.2, --operation mean, 5 kb resolution); clones were not merged at this stage. For each bigWig, chrX was binned at 5 kb using bioframe v0.8.0, and a two-state Gaussian HMM (hmmlearn v0.3.3, GaussianHMM; n_components = 2, covariance_type = “diag”, n_iter = 15, tol = 0.01, min_covar = 0.001, random_state = 42; https://github.com/hmmlearn/hmmlearn) was fitted to the binned signal. The state with the lower emission mean was assigned as “valley”; the higher-mean state as “elevation”. To minimise the influence of mappability artefacts on model fitting, ENCODE mm10 blacklist v2 regions were excluded from chrX, and only contiguous non-blacklisted intervals ≥ 500 kb were used for training. Valley intervals were exported as BED and bigWig files for downstream analyses. To test for the presence of CTCF at H3K27me3 valley boundaries, we defined a window extending 10 kb outward and 50 kb inward from each boundary. This asymmetric window was chosen because CTCF binding was expected to occur outside H3K27me3-covered regions, while allowing for potential uncertainty in the precise boundary location.

#### Statistics and reproducibility

No statistical methods were used to predetermine sample size. The experiments were not randomized and the investigators were not blinded to allocation during experiments and outcome assessment. The statistical tests used, corresponding p-values, and sample sizes are indicated in the figure panels, figure legends, or methods.

## Data visualization

Unless stated otherwise, box plots show the median (center line), first and third quartiles (box limits), and whiskers extending to the most extreme values within 1.5× the interquartile range and bars in bar charts represent the mean and error bars indicate standard deviation (SD).

## Data availability

RNA-seq, Capture Hi-C and CUT&RUN data generated in this study are deposited in the Gene Expression Omnibus under accession number <u>GSE330701</u> and accession password: XXXXXX. This study uses previously published datasets available under GSE259399 (CL30.7 Ctrl Mecp2 (GSM8147323), E6 Ctrl Mecp2 (GSM8147326) and E6 Ctrl Kdm5c (GSM8147330)). All sequencing datasets were aligned using NCBI RefSeq GRCm38 genome assembly and using the corresponding RefSeq transcript annotations Mus_musculus.GRCm38.100.gtf.

## Code availability

The preprocessing workflows for RNA-seq and CUT&RUN data are available via GitHub at https://github.com/yuviaapr/allele-specific_RNA-seq and https://github.com/yuviaapr/allele-specific_CUTandRUN. The code used to analyse RNA-seq and CUT&RUN allelic data is based on code available via GitHub at https://github.com/odomlab2/xist_project. The pipelines for CUTandRUN/RNA-Seq normalisation, H3K27me3 valley identification and analysis, loops/compartments calling and METALoci analysis are available via GitHub at https://github.com/encent/xci-pipelines. METALoci code is available via GitHub at https://github.com/3DGenomes/METALoci.

## Acknowledgements

This work was funded by an ERC Advanced Investigator award to E.H. (XPRESS - award no. AdG671027) and the European ITN Innovative and Interdisciplinary Network ‘ChromDesign’, under the Marie Skłodowska-Curie grant agreement no. 813327 to E.H and M.A.M-R. This project has received funding from the European Union’s Horizon 2020 research and innovation programme under the Marie Skłodowska-Curie grant agreements No 882771 to Y.A.P.-R. and No 838408 to A.L. M.A.M-R also acknowledges support from the Spanish Ministerio de Ciencia, Innovacion y Universidades grant number PID2023-151484NB-I00.

## Author contributions

A.L. and E.H. conceived the study. M.A.M-R. conceived and designed the computational framework to analyse the 3D (Hi-C) data and integrate it with epigenomic data. A.L. generated the WT NPC clones with C.P., set up and performed the Capture Hi-C experiments in WT clones, performed some of the RNA-seq, CUT&RUN and CRISPR-Cas9 experiments. A.H. performed all Capture Hi-C, RNA-seq, CUT&RUN, CRISPR-Cas9 experiments trained by A.L, with technical support for some cell line generation, sample preparation and processing from I.R, B.K. and C.S., A.H. pre-processed the RNA-seq and CUT&RUN data and performed downstream analysis of these datasets unless stated otherwise. N.B. performed inter-clonal normalization of CUT&RUN data, H3K27me3 valley calling and analysis, loop and compartment analysis of Capture Hi-C and METALoci analysis. Y.A.P.-R. developed the processing RNA-seq and CUT&RUN pipelines, generated and processed some of the CUT&RUN and RNA-seq datasets with help from L.C. and performed differential expression analysis of RNA-seq data. N.S. processed the Capture Hi-C with A.H. T.P. generated the CTCF degron targeting vector and participated in generating the RAD21-dTAG ESCs. L.V. prepared RNA-seq and Capture Hi-C libraries M.A.M-R. supervised N.B. for computational analyses and provided input to the whole study. A.L., E.H. and M.A.M-R. supervised the work and wrote the manuscript with input from A.H. and N.B.

## Notes

### Competing Interest Statement

The authors have declared no competing interest.

