## Extended Data Figures for "X-inactivation escapee domains are CTCF-cohesin independent chromatin compartments"

### Extended Data Figure 1

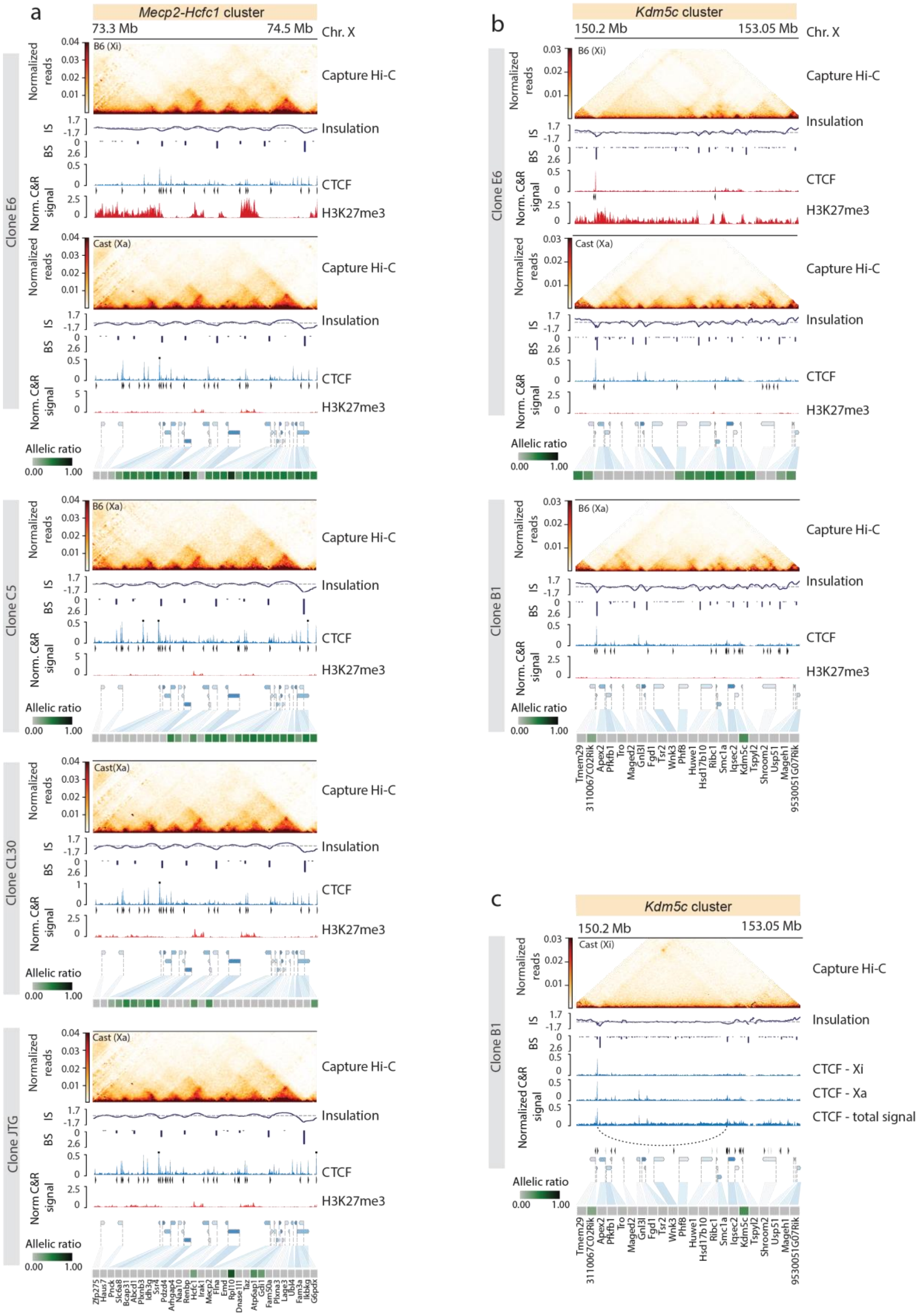

**Extended Data Figure 1**

**a**, Capture Hi-C interaction maps, insulation scores, allele-specific CTCF CUT&RUN profiles and called peaks, and H3K27me3 CUT&RUN profiles across the *Mecp2–Hcfc1* escape cluster are shown for NPC clones E6, C5, CL30 and JTG (corresponding to the Xa chromosomes of the clones shown in Fig. 1c), and for both the Xi and Xa of clone E6. Capture Hi-C data are shown at 10 kb resolution. CTCF motif orientation is indicated by arrowheads. Heatmaps of allelic ratios of genes within the cluster are shown. **b**, As in **a**, for the *Kdm5c* cluster. **c**, As in **b**, for the Xi of NPC clone B1, highlighting a prominent long-range looping interaction (dashed arc) and corresponding CTCF binding profiles on the Xi, Xa, and total signal. Enrichment of CTCF in close proximity upstream of the loop anchor is detectable only in the total signal due to the absence of informative SNPs at this locus. [IS - insulation score; BS - boundary score]

### Extended Data Figure 2

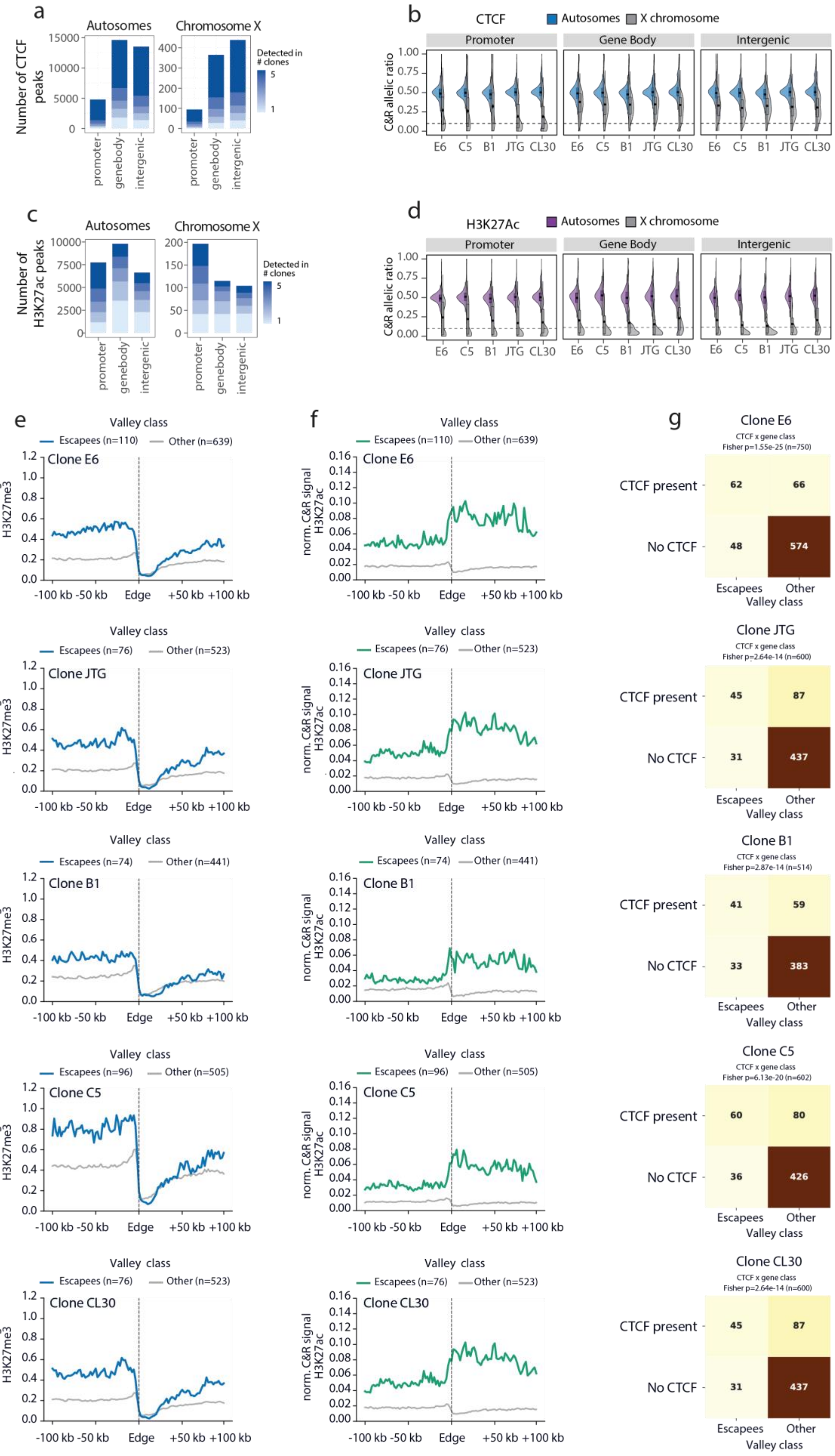

**Extended Data Figure 2**

**a**, Number of CTCF peaks detected at promoters, gene bodies, and intergenic regions on autosomes (left) and the X chromosome (right). Bars are stacked and colored by the number of NPC clones (1–5, B1, C5, JTG, E6 and CL30) in which each peak was detected, with darker shading indicating peaks shared across a greater number of clones. **b**, Violin plots showing the distribution of allelic ratios (Xi/(Xi+Xa)) for CTCF peaks across promoters, gene bodies and intergenic regions on autosomes (blue) and the X chromosome (grey). Autosomal peaks are predominantly biallelic; X-linked peaks show enrichment on the Xa relative to the Xi (allelic ratio < 0.5). **c-d**, As in **a-b** for H3K27ac (autosomes:violet and X chromosome:grey). **e**, Mean H3K27me3 signal in 100 kb windows centred on the edges of H3K27me3 valleys containing escapees (blue) or other H3K27me3 valleys (grey; valleys overlapping silent genes, not expressed genes or intergenic region; see Fig. 2g) on the Xi of NPC clones E6, JTG, B1, C5 and CL30, calculated from normalised, replicate-merged bigWig tracks at 5 kb resolution; n, edges of valleys per class. **f**, As in **e** for H3K27ac (escapees: green; other: grey). **g**, 2 × 2 contingency table of H3K27me3 valley edges stratified by class (escapees versus other, see Fig. 2g) and CTCF presence (CTCF present versus No CTCF). A valley edge was scored as CTCF present if a clone-specific CTCF CUT&RUN peak fell within 50 kb inward and 10 kb outward of the valley edge. Cell counts and a two-sided Fisher's exact test p-value are shown (n, number of valley edges per clone).

**Extended Data Figure 3**

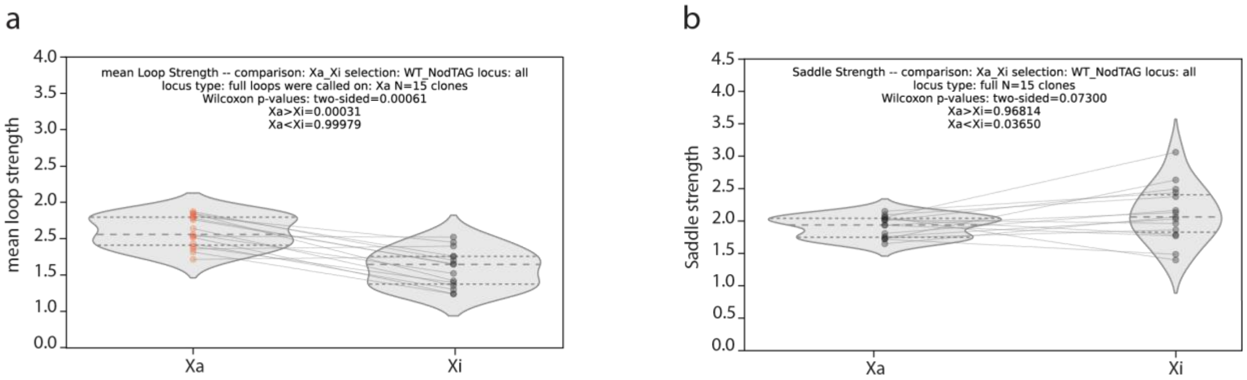


**Extended Data Figure 3**

**a**, Mean loop strength per clone on Xa versus Xi (at genomic locations of loops called on Xa), pooling both *Mecp2-Hcfc1* and *Kdm5c* clusters (N = 15; WT NPC clones and dTAG clones in untreated conditions, see methods). For each clone, the mean observed-over-expected contact frequency across all consensus loops was computed; Xa and Xi values of the same clone are connected by a line. Inset: two-sided Wilcoxon signed-rank p-value and corresponding one-sided p-values. **b**, As in **a** for compartment strength, defined as (AA + BB) / (AB + BA) from the cooltools saddle (N = 38 bins, qrange = (0.025, 0.975)). Compartment strength was computed independently on the Xa and the Xi map of each clone, yielding paired Xa/Xi measurements (N = 15; WT NPC clones and dTAG clones in untreated conditions, see methods).

**Extended Data Figure 4**

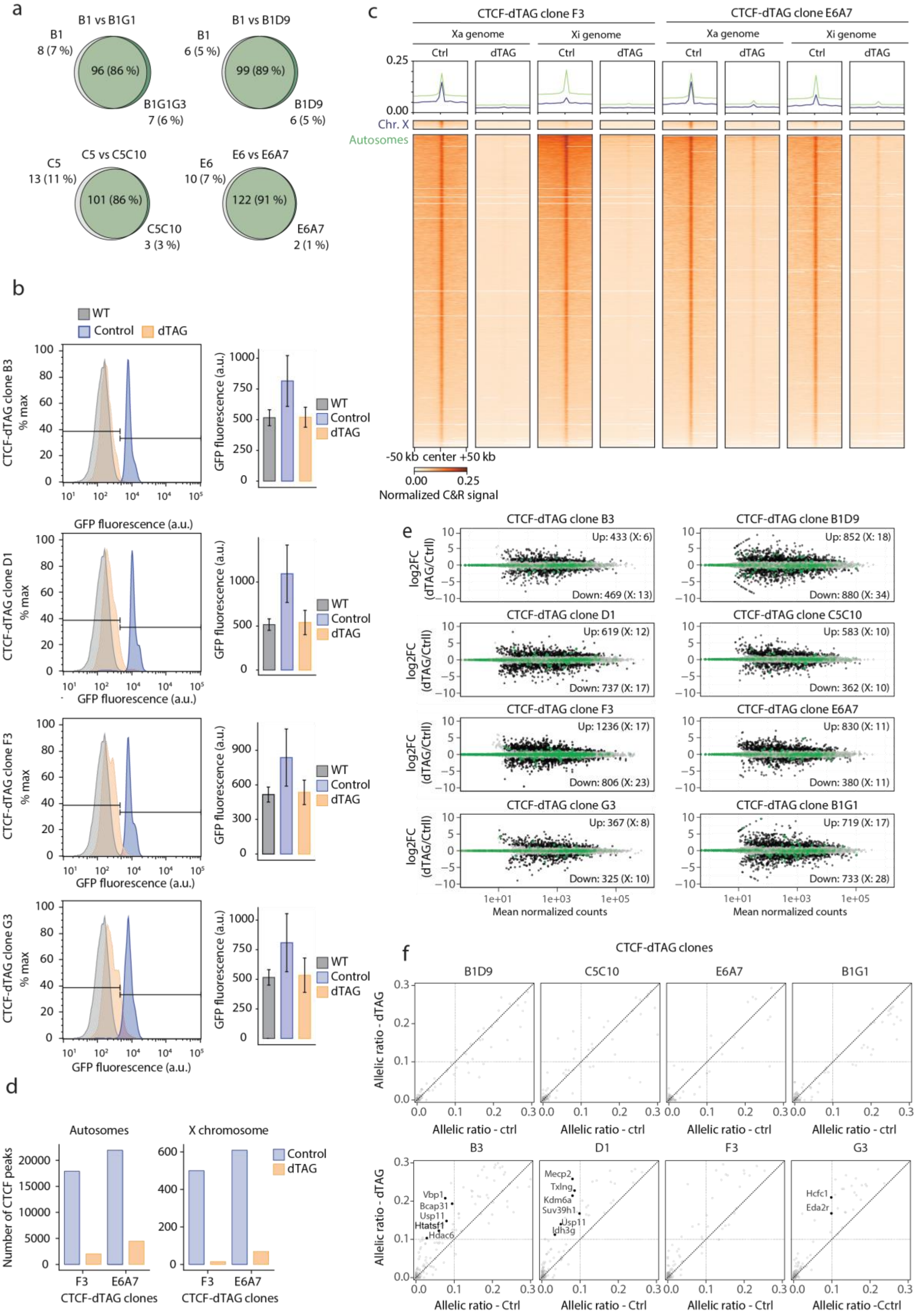

**Extended Data Figure 4**

**a**, Venn diagrams and Jaccard indices showing the overlap of escapees between CTCF-dTAG NPC clones and their corresponding parental wild-type clones. Only genes detected in both datasets were included in the analysis. **b**, Flow cytometric validation of acute CTCF depletion following 30 min of dTAG treatment in CTCF-dTAG NPC clones B3, D1, F3 and G3. Left, histograms showing GFP fluorescence intensity distributions in wild-type, untreated CTCF-dTAG, and dTAG-treated NPCs. Right, quantification of mean GFP fluorescence intensity across live cells. Bars indicate mean values and error bars represent standard deviation. **c**, Heatmaps and aggregate profiles showing CTCF CUT&RUN signal across all chromosomes, with allelic signals assigned to the haplotypes carrying the Xa and Xi, before and after dTAG treatment in CTCF-dTAG NPC clones F3 (left) and E6A7 (right). Signal is shown across all CTCF peaks located on the X chromosome (n = 1179, blue) and autosomes (n = 42933, green), including ±50 kb flanking regions. **d**, Number of CTCF peaks detected on autosomes and the X chromosome in untreated and dTAG-treated CTCF-dTAG clones E6A7 and F3. **e**, MA plots for CTCF-dTAG clones showing log2 fold change (dTAG/control) as a function of mean normalized counts. Autosomal genes are shown in grey (non-significant) and black (significant (abs. log2FC > 1; adjusted $P < 0.05$)); X-linked genes are shown in light green (non-significant) and dark green (significant (abs. log2FC > 1; adjusted $P < 0.05$)). **f**, Boxplots showing allelic ratios of X-linked genes in CTCF-dTAG clones in control and dTAG-treated conditions. Statistical significance was assessed using paired two-sided Wilcoxon signed-rank tests with Benjamini–Hochberg correction for multiple testing. Twelve of 361 silent genes across eight clones show a significant increase of allelic ratio and pass the escapee threshold (allelic ratio > 0.1) upon dTAG treatment. These genes are highlighted by black dots and labelled with their gene names.

#### Extended Data Figure 5

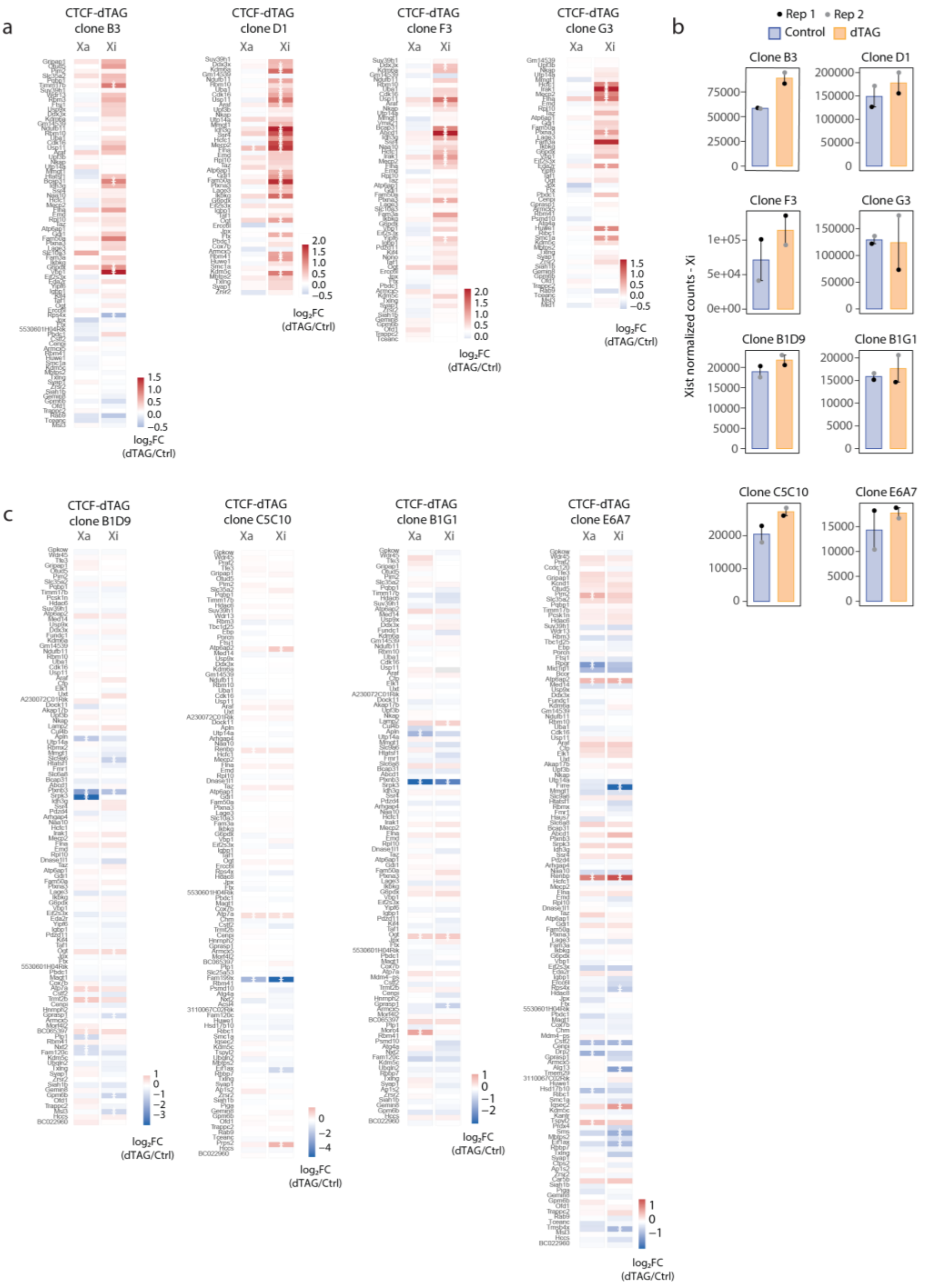


#### Extended Data Figure 5

**a**, Heatmaps of allele-specific log 2 fold changes (Log2FC) in gene expression for escapees following 4 days of dTAG treatment in four CTCF-dTAG clones derived from CRISPR-Cas9-targeted mESCs. Data for X-linked genes with allelic ratio > 0.1 in either the control or dTAG-treated condition is shown. Values are shown separately for the Xa and Xi. Genes are ordered by chromosomal position. White asterisks indicate significantly differentially expressed genes (adjusted $P < 0.05$). **b**, Xist expression levels across eight CTCF-dTAG NPC clones in untreated and 4-day dTAG-treated conditions. Data are shown for two biological replicates. **c**, As in **a**, but for four CTCF-dTAG NPC clones derived by direct CRISPR-Cas9 targeting in wild-type NPCs.

**Extended Data Figure 6**

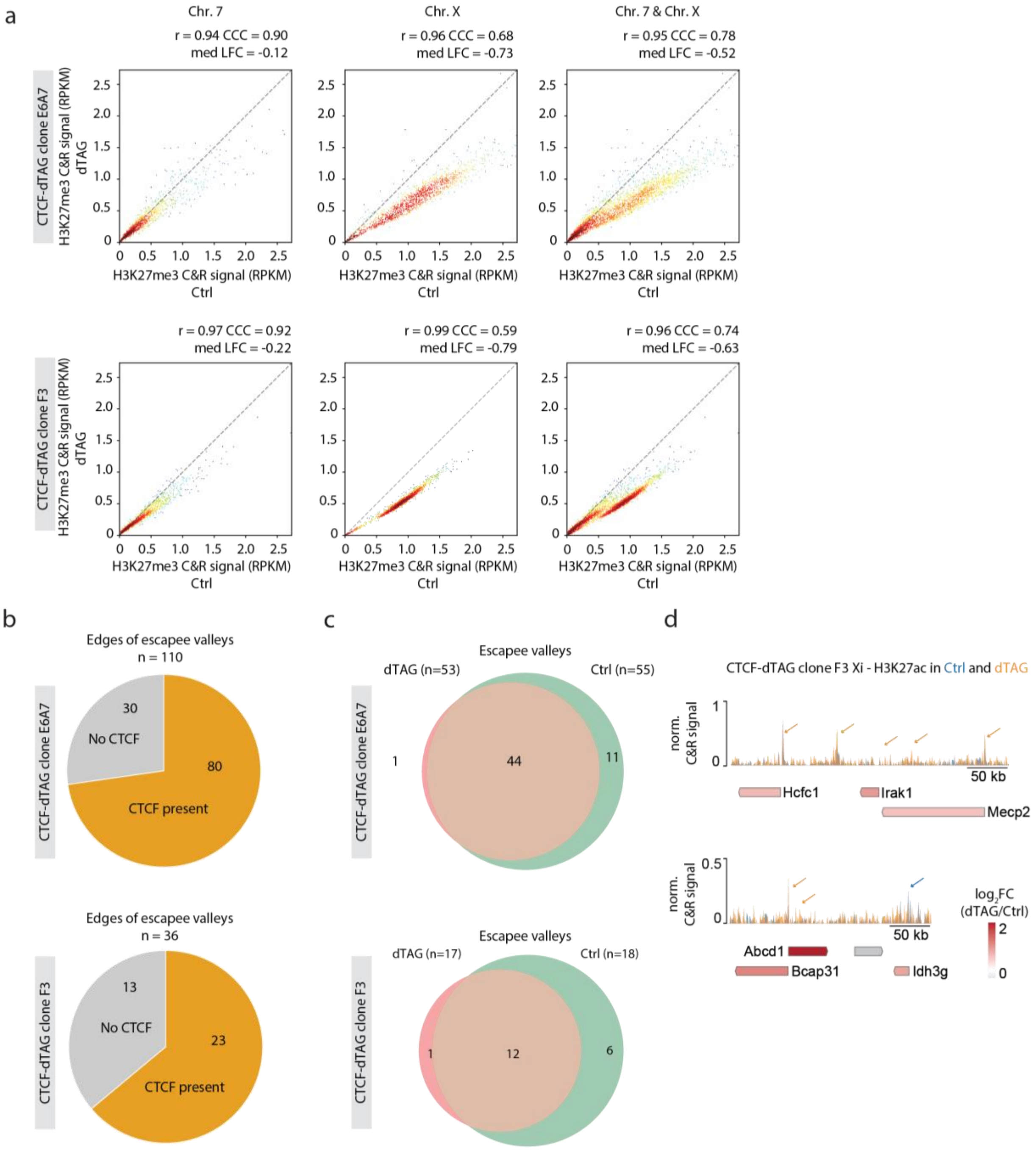

**Extended Data Figure 6**

**a**, Scatter plots comparing H3K27me3 coverage on chromosome 7 (left), chromosome X (middle) and both (right) in control versus dTAG-treated conditions for CTCF-dTAG clones E6A7 and F3. Points are colored by local density (red/yellow = high density, blue = low density), with the dashed gray line indicating the diagonal (Ctrl = dTAG, i.e., no change). Pearson correlation (r), concordance correlation coefficient (CCC), and median log2 fold-change (med LFC) are shown above each plot. **b**, Pie charts showing the presence or absence of CTCF peaks at the edges of H3K27me3 valleys encompassing at least one escapee in CTCF-dTAG clones E6A7 and F3. Slices indicate the proportion of edges with a CTCF peak (orange) within ≤50 kb inward and ≤10 kb outward of the valley edge, or without a nearby CTCF peak (grey). *n*, valley edges per clone. **c**, Venn diagrams comparing H3K27me3 valleys on the Xi overlapping at least one escapee between dTAG (red) and control (green) conditions, for CTCF-dTAG clones E6A7 and F3. Valleys were identified as in Fig. 2i. Overlap is assessed region-by-region with bioframe.overlap under the NodTAG-centric convention (Ctrl ∩ dTAG = Ctrl valleys overlapping at least one dTAG valley). **d**, H3K27ac CUT&RUN signal across two regions of the *Mecp2-Hcfc1* cluster encompassing genes with significantly increased Xi expression following CTCF depletion in CTCF-dTAG clone F3. Profiles from control (blue) and dTAG-treated (orange) cells are overlaid. Blue and orange arrows indicate H3K27ac peaks with higher signal in control and dTAG-treated cells, respectively. Genes are shown below the tracks. Genes with significantly increased Xi expression following CTCF depletion are labeled and gene symbols are colored according to the Xi log2FC(dTAG/Ctrl).

**Extended Data Figure 7**

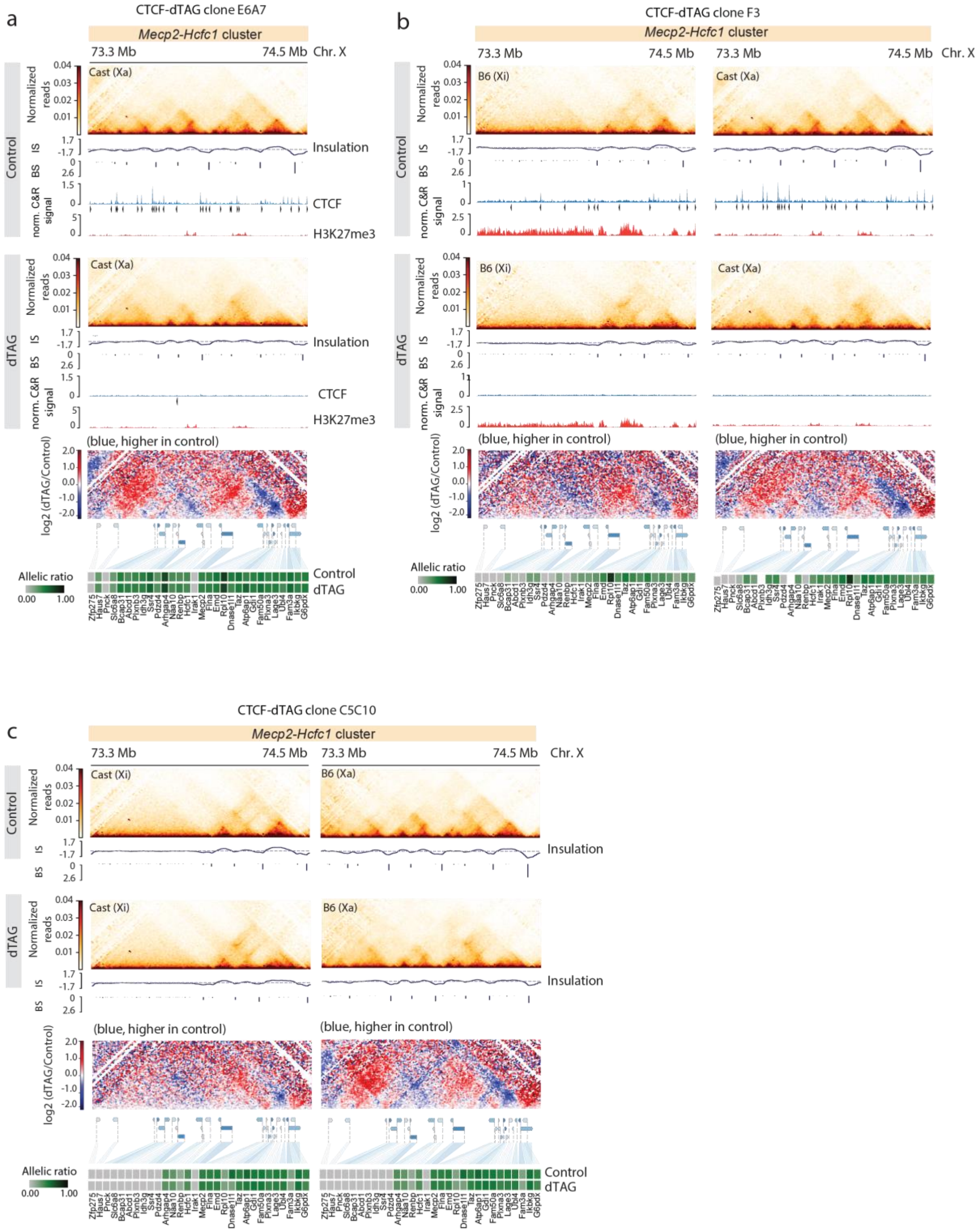

**Extended Data Figure 7**

**a**, Capture Hi-C interaction maps, insulation scores, allele-specific CTCF CUT&RUN profiles and called peaks, and H3K27me3 CUT&RUN profiles across the *Mecp2–Hcfc1* cluster. Data is shown for the Xa in CTCF-dTAG NPC clone E6A7 under control and dTAG-treated conditions. Capture Hi-C data is shown at 10 kb resolution. CTCF motif orientation is indicated by arrowheads. Differential Capture Hi-C maps show changes in chromatin interactions between control and 4-days dTAG-treated conditions. Heatmaps of allelic ratios for all genes within the cluster are shown for the control and dTAG-treated conditions. **b**, As in **a**, but for the Xa and Xi of CTCF-dTAG NPC clone F3 under control and dTAG-treated conditions. **c**, Capture Hi-C interaction maps and insulation scores across the *Mecp2–Hcfc1* cluster on the Xa and Xi of CTCF-dTAG NPC clone C5C10 under control and dTAG-treated conditions. Capture Hi-C data is shown at 10 kb resolution. Differential Capture Hi-C maps show changes in chromatin interactions between control and 4-days dTAG-treated conditions. Heatmaps of allelic ratios for all genes within the cluster are shown for the control and dTAG-treated conditions. [IS - insulation score; BS - boundary score]

#### Extended Data Figure 8

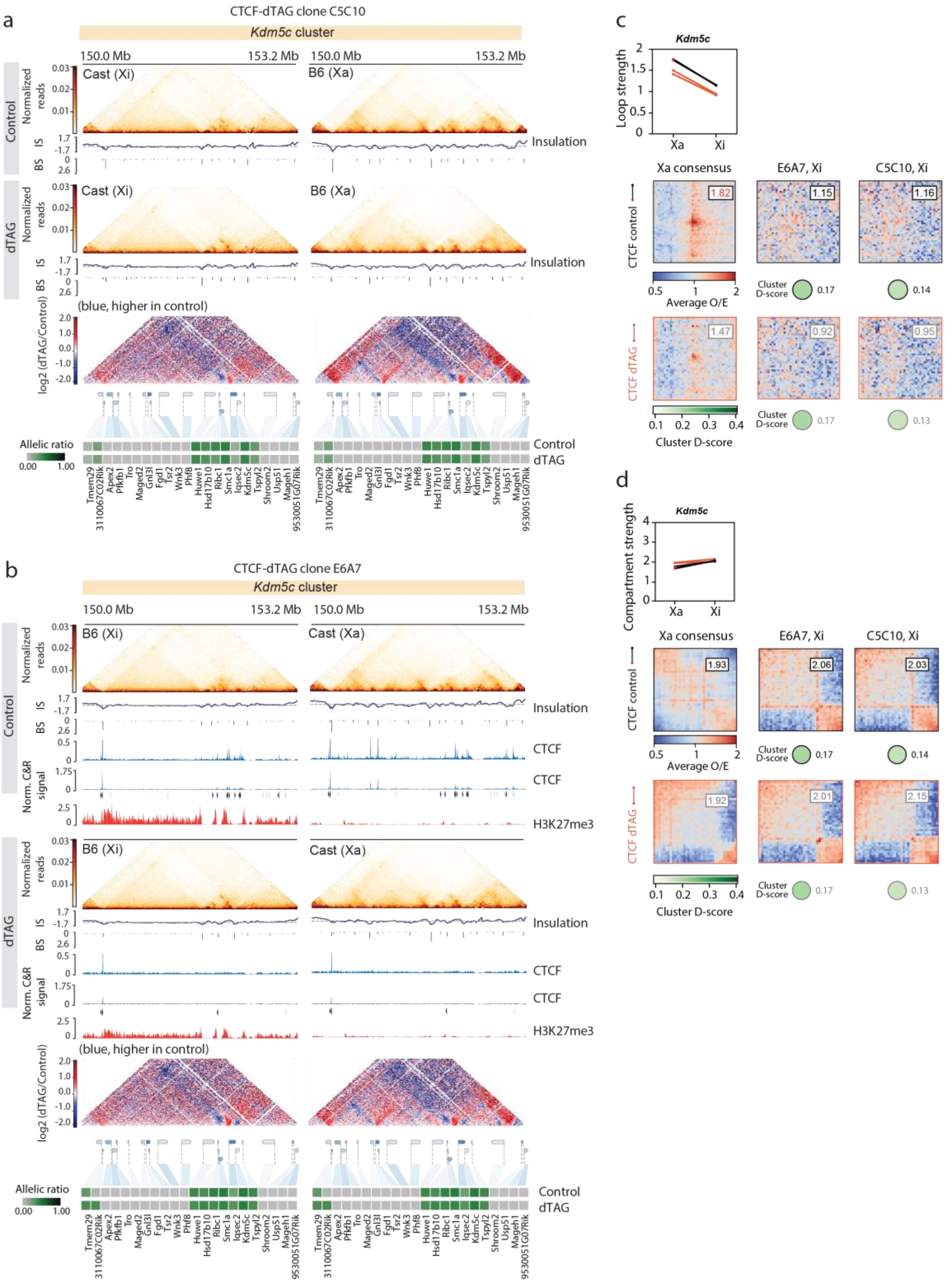

**Extended Data Figure 8**

**a**, Capture Hi-C interaction maps and insulation scores across the *Kdm5c* cluster on the Xa and Xi of CTCF-dTAG NPC clone C5C10 under control and dTAG-treated conditions. Capture Hi-C data is shown at 10 kb resolution. Differential Capture Hi-C maps show changes in chromatin interactions between control and 4-days dTAG-treated conditions. Heatmaps of allelic ratios for all genes within the cluster are shown for the control and dTAG-treated conditions. **b**, Capture Hi-C interaction maps, insulation scores, allele-specific CTCF CUT&RUN profiles and called peaks, and H3K27me3 CUT&RUN profiles across the *Kdm5c* cluster. Data is shown for the Xa and Xi in CTCF-dTAG NPC clone E6A7 under control and dTAG-treated conditions. Capture Hi-C data is shown at 10 kb resolution. CTCF motif orientation is indicated by arrowheads. Differential Capture Hi-C maps show changes in chromatin interactions between control and 4-days dTAG-treated conditions. Heatmaps of allelic ratios for all genes within the cluster are shown for the control and dTAG-treated conditions. **c**, Loop strength at the *Kdm5c* cluster across CTCF-dTAG clones E6A7 and C5C10, for the control (middle) and dTAG-treated (bottom) conditions. Per-clone mean loop strength on Xa and Xi (top; one line per clone – control (black) and dTAG-treated (orange) conditions; loops were called on Xa), together with average observed-over-expected loop pile-up matrices at 5 kb resolution centred on consensus *Kdm5c* loops (see methods); mean O/E in the central 3 × 3 pixels is shown at top right of each pile-up. The Xa consensus pile-up uses the merged Xa cooler (see methods); Xi pile-ups are clone-specific (E6A7, C5C10). Green circles below each pile-up reflect the per-clone average cluster D-score. **d**, As in **c** for compartment strength. Saddle plots are shown for the Xa consensus (see methods) and clone-specific Xi chromosomes. [IS - insulation score; BS - boundary score]

#### Extended Data Figure 9

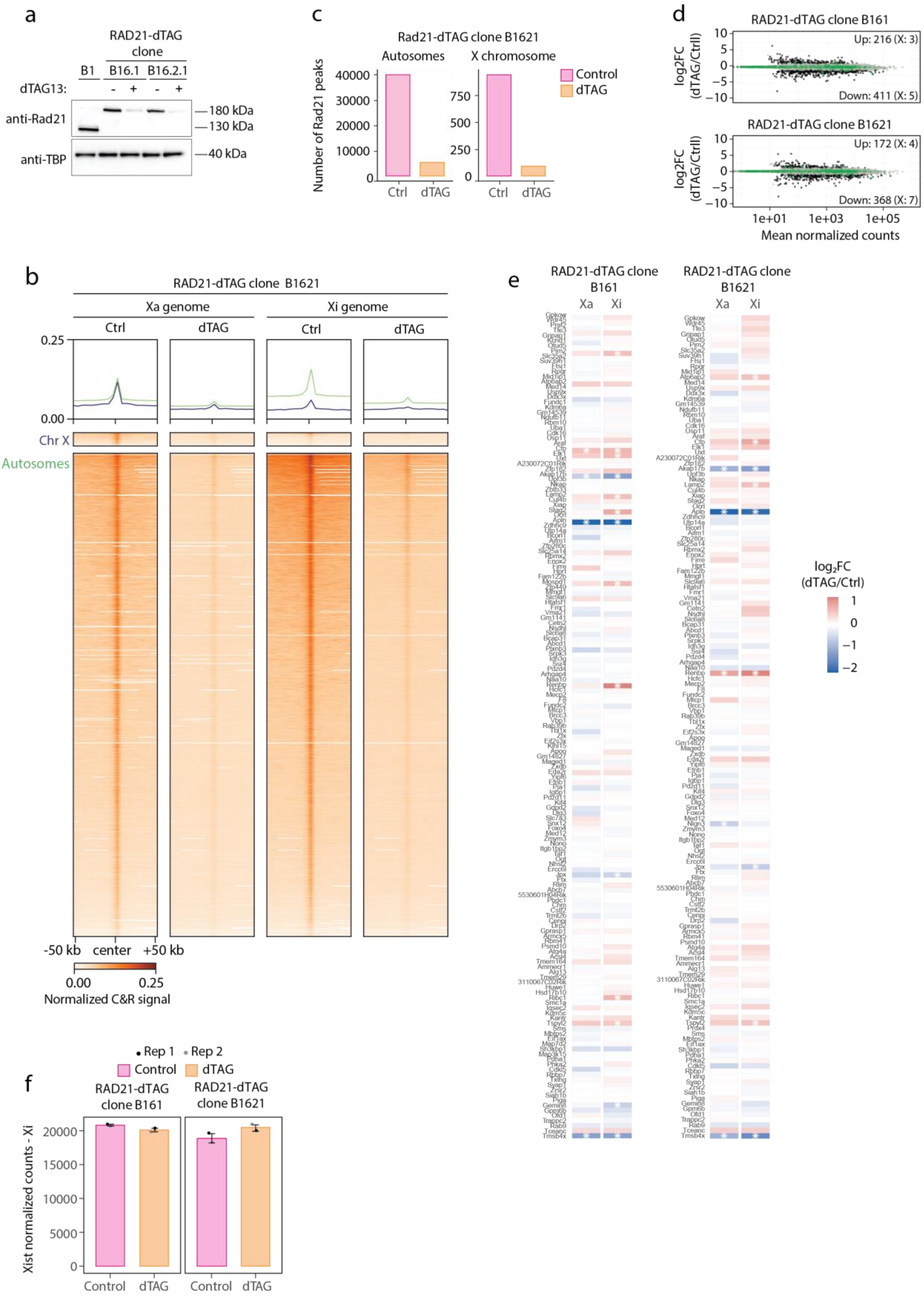

**Extended Data Figure 9**

**a**, Western blot showing RAD21 protein levels in two independent RAD21-dTAG NPC clones (B161 and B1621) treated (+) or untreated (−) with dTAG. Wild-type NPC clone B1 is shown for comparison. TBP served as the loading control. **b**, Heatmaps and aggregate profiles showing RAD21 CUT&RUN signal across all chromosomes, with allelic signals assigned to the haplotypes carrying on the Xa and Xi, before and after dTAG treatment before and after dTAG treatment in RAD21-degron NPC clone B1621. Signal is shown across all RAD21 peaks located on the X chromosome (n = 1213, blue) and autosomes (n = 44343, green), including ±50 kb flanking regions. **c**, Number of RAD21 peaks detected on autosomes and the X chromosome in untreated and dTAG-treated RAD21-dTAG NPC clone B1621. **d**, MA plots for RAD21-dTAG NPC clone B1621 showing log2 fold change (dTAG/control) as a function of mean normalized counts. Autosomal genes are shown in grey (non-significant) and black (significant (abs. log2FC > 1; adjusted P < 0.05)); X-linked genes are shown in light green (non-significant) and dark green (significant (abs. log2FC > 1; adjusted P < 0.05)). **e**, Heatmaps of allele-specific log2 fold changes (Log2FC) changes in gene expression for escapees following 6 hours of dTAG treatment in two RAD21-dTAG NPC clones derived by differentiation of CRISPR-Cas9-targeted mESCS. Data for X-linked genes with allelic ratio > 0.1 in either the control or dTAG-treated condition is shown. Log2 fold changes (log2FC) values are shown separately for the Xa and Xi. Genes are ordered by chromosomal position. White asterisks indicate significantly differentially expressed genes (adjusted $P < 0.05$). **f**, Xist expression levels across RAD21-dTAG NPC clones B161 and B1621 in untreated and 6-hour dTAG-treated conditions. Data are shown for two biological replicates.

#### Extended Data Figure 10

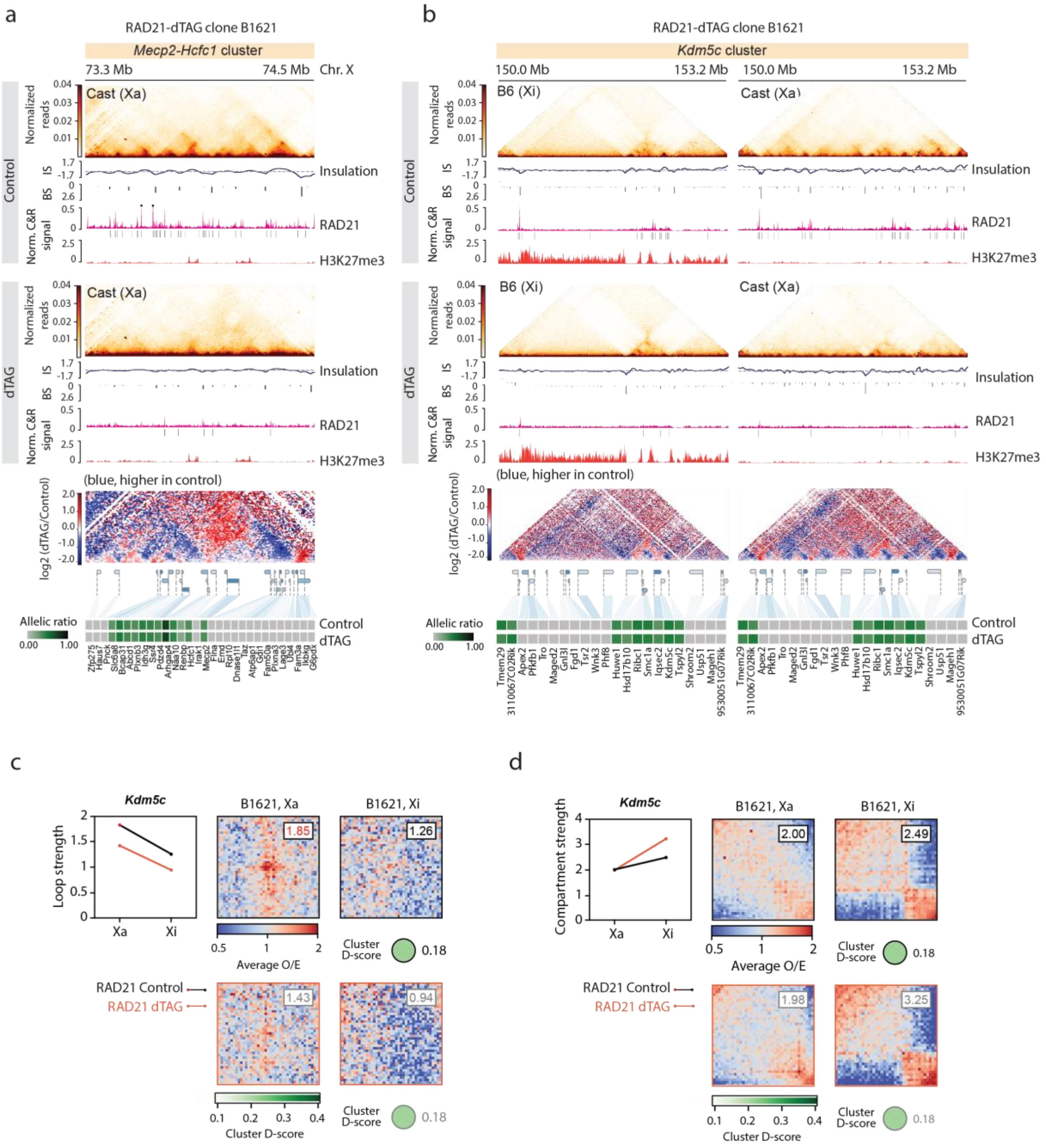

**Extended Data Figure 10**

**a,** Capture Hi-C interaction maps, insulation scores, allele-specific RAD21 CUT&RUN profiles and called peaks, and H3K27me3 CUT&RUN profiles across the *Mecp2–Hcfc1* cluster. Data is shown for the Xa in RAD21-dTAG NPC clone B16121 under control and dTAG-treated conditions. Capture Hi-C data is shown at 10 kb resolution. Differential Capture Hi-C maps show changes in chromatin interactions between control and 6-hour dTAG-treated conditions. Heatmaps of allelic ratios for all genes within the cluster are shown for the control and dTAG-treated conditions. **b**, As in **a**, but for the *Kdm5c* cluster on the Xa and Xi of RAD21-dTAG NPC clone B16121 under control and dTAG-treated conditions. **c**, Loop strength at the *Kdm5c* cluster in RAD21-dTAG clone B1621, for the control (top) and dTAG-treated (bottom) conditions. Per-clone mean loop strength on Xa and Xi (top-left; one line per clone – control (black) and dTAG-treated (orange) conditions; loops were called on Xa); together with average observed-over-expected loop pile-up matrices at 5 kb resolution centered on consensus *Kdm5c* loops (see methods); mean O/E in the central 3 × 3 pixels is shown at top right of each pile-up. Both Xa and Xi pile-ups are clone-specific. Green circles below each pile-up reflect the per-clone average cluster D-score. **d,** As in **c**, for compartment strength. Saddle plots are shown for the Xa consensus (see methods) and clone-specific Xi chromosomes. [IS - insulation score; BS - boundary score]
